# Fitness benefit plays a vital role in the retention of the *Pi-ta* susceptible alleles

**DOI:** 10.1101/2020.07.30.229831

**Authors:** Jia Liu, Yue Wang, Suobing Zhang, Fan Wu, Long Wang, Jiayu Xue, Yanmei Zhang, Pengfei Xie, Yueyu Hang, Xiaoqin Sun

## Abstract

In plants, large numbers of *R* genes, which segregate as loci with alternative alleles conferring different resistance to pathogens, have been maintained over a long evolutionary time. In theory, there seem to be no reason for hosts to harbor these susceptible alleles in view of their null contribution to resistance. As such, why should populations support disease-susceptible individuals along with disease-resistant individuals? In rice, a single copy *R* gene *Pi-ta* segregates for two expressed clades of alleles, one resistant and the other susceptible. We discovered that knockout of the *Pi-ta* susceptible alleles induced drastic fitness decline in the absence of pathogens. Gene expression profiling and endogenous hormones quantification showed that the susceptible alleles might serve as an off-switch to the downstream immune signaling, thus contributing to fine-tuning of plant defense response. The fitness benefit of carrying a susceptible *Pi-ta* allele provides a plausible explanation for their retention in the genome.

## Introduction

In plants, defense pathways against microbial and fungal pathogens are triggered in plants by recognition and signal transduction system encoded by large diverse multigene families, called *R-*genes (Jones and Dangl 2006). Although genetic resistance provides an effective method for the control of plant diseases, mounting a defense response frequently comes with the cost of reduction in growth and reproduction, carrying critical implications for natural and agricultural plant populations (Brown 2002). Research on fitness effects was pioneered by Van der Plank (1963) in his study of the resistance of potato to late blight (*Phytophthora infestans*) and has been documented in other plants. For example, the wheat (*Triticum aestivum*) streak mosaic virus *R* gene *Wsm1* is associated with a mean yield reduction of 21% (Sharp, et al. 2002), the wheat stem rust *R* gene *Sr26* has a 9% yield penalty (Brown 2002), and the barley (*Hordem vulgare*) *mlo R* gene has a 4.2% yield penalty (Jørgensen 1992). Research on fitness effects of resistance is currently enjoying a surge of interest, especially in studying the molecular mechanisms underlying the trades-offs between yield and disease resistance and proposing new breeding strategies for selecting high-yield cultivars with minimum fitness costs in crops. For example, *Pigm* and *Xa4* are discovered to confer disease resistance to the rice blast fungus (*Magnaporthe oryzae*) and the rice bacterial blight (*Xanthomonas oryzae* pv. *oryzae*), respectively, both without compromising grain yield (Deng, et al. 2017; Hu, et al. 2017).

*R* genes can evolve rapidly, presumably to keep pace with the evolution of pathogens, otherwise pathogens may evolve out of the detection scope. Molecular genetic studies of plant *R* genes have revealed extensive variation in *R* gene loci (Bakker, et al. 2006), with fitness effects acting as one of the key selective drivers of *R* gene evolution. Mathematical models generally suggest that a trade-off between costs of resistance in pathogen-free environments and benefits of resistance under infection is required to explain the long-lived presence/absence polymorphisms of several *R* genes (Stahl, et al. 1999; Bergelson, et al. 2001). Indeed, using isolines, a high cost of resistance measured in the absence of disease has been determined for two *R* genes, *Rps5* and *Rpm1*, in *Arabidopsis thaliana* (Tian, et al. 2003; Karasov, et al. 2014). Both *Rps5* and *Rpm1* exist in nature as a long-lived, presence/absence polymorphism for the entire gene for resistance (R) and susceptibility (S), respectively. In both cases, resistant isolines suffer a 5-10% fitness cost relative to null isolines in the absence of disease. Interestingly, our previous fitness trials testing the consequences of the *Rps2* gene (Macqueen, et al. 2016), which exists as an ancient balanced polymorphism with two long-lived clades of alleles, one resistant and the other susceptible to *Pseudomonas syringae* pv. *avrRpt2*, found no fitness cost for encoding the resistance allele in the absence of infection. So it’s hypothesized that loci which exhibit presence/absence polymorphisms are most likely to have large fitness costs, while those loci that exhibit selection on multiple alleles may have negligible costs. Such a hypothesis seeks to explain how host genomes can tolerate the possible genetic load associated with a vast repertoire of *R* genes.

Rice blast disease is one of the most threatening diseases for rice production worldwide. The *Pi-ta* gene in rice has been effectively used to fight against blast disease globally (Bryan, et al. 2000; Jia, et al. 2000). *Pi-ta* is a single copy gene that encodes a predicted cytoplasmic protein with a nucleotide binding site (NBS) and leucine-rich repeats (LRR) (Bryan, et al. 2000; Jia, et al. 2000). Interestingly, *Pi-ta* is present in every rice accession studied to date, none of which has missing data or deletions called for *Pi-ta* (Jia, et al. 2016). This gene has been shown to be segregated for both resistant and susceptible alleles to *Magnaporthe oryzae* (AVR-*Pita*) (Bryan, et al. 2000; Jia, et al. 2000). The resistant *Pi-ta* alleles might have recently arisen with nearly no variation and a low occurrence rate, during a blast pathogen-host shift from Italian millet to rice (Huang, et al. 2008; Wang, et al. 2008). On the contrary, most genetic variations occur in susceptible *Pi-ta* alleles, suggestive of a relaxed positive selection (Lee, et al. 2009; Thakur, et al. 2013). The undeniable benefits of resistance should have driven the resistant alleles to fixation, while in theory there would thus be no reason for hosts to harbor susceptible alleles. Why should rice populations still retain disease-susceptible individuals along with disease-resistant individuals? We hypothesized that the *Pi-ta* susceptible allele must have another beneficial function to permit its retention. Wang et al. (2015) reported that the *Pi-ta* susceptible alleles were associated with heavier seed weight based on genome-wide association analysis (GWAS), indicative of the fitness effect of the susceptible alleles on yield. However, it is still unclear if the effect of *Pi-ta* on yield is due to a direct effect of the gene itself or another yield-related gene located in the same genomic region. After all, genes that are linked to one *R* gene may also affect yield (Brown 2002).

Empirical evidence for the fitness effects of *R* genes is still controversial, because some *R* genes may be tightly linked to other yield-related genes (Ortelli, et al. 1996) and because fitness effects are affected by environmental conditions and may vary by evaluation method (Laine 2016). Therefore, pairs of transgenic lines with rigorously controlled background and field trials should be the best materials for studying the fitness effects of individual *R* genes (Ning, et al. 2017).

Herein, to discover the role of fitness effects on the retention of the *Pi-ta* susceptible allele, we simulated a loss of function of the susceptible *Pi-ta* allele to detect any subsequent fitness changes. Loss of function of the susceptible *Pi-ta* allele was induced by targeted mutation using the CRISPR/Cas9 system. In particular, we screened a rigorously controlled line without mutations in siblings of the knockout mutants as the control. We measured and compared the fitness in isogenic lines with and without a functional *Pi-ta* allele in the absence of disease. We additionally verified the robustness of our results by using five independent isogenic lines for the mutation of *Pi-ta*. Our results revealed that the knockout of the *Pi-ta* susceptible *alleles* was costly in terms of fitness. Gene expression profiling of these isogenic lines showed that the *Pi-ta* susceptible alleles contributed to fine-tuning of the downstream defense response, explaining their retention in the genome.

## Results

### Generation of isogenic lines of the *Pi-ta* gene using CRISPR/Cas9

*Pi-ta* is a single copy gene that encodes a predicted cytoplasmic protein with NBS and LRR domains (Bryan, et al. 2000; Jia, et al. 2000). Nipponbare (*Oryza sativa* L. ssp. *japonica*), which served as the genetic background in this study, possesses a susceptible allele of the *Pi-ta* gene in its genome. Two target sites for CRISPR/Cas9 editing were located in proximity to the 5’ end and the 3’ end of the gene (Figure 1A). Synthesized oligos were inserted into the CRISPR/Cas9 binary vector for editing (Figure 1B). Subsequently, the two constructed vectors were transformed into Nipponbare using the Agrobacterium-mediated method (Hiei, et al. 1994).

**Figure 1.**
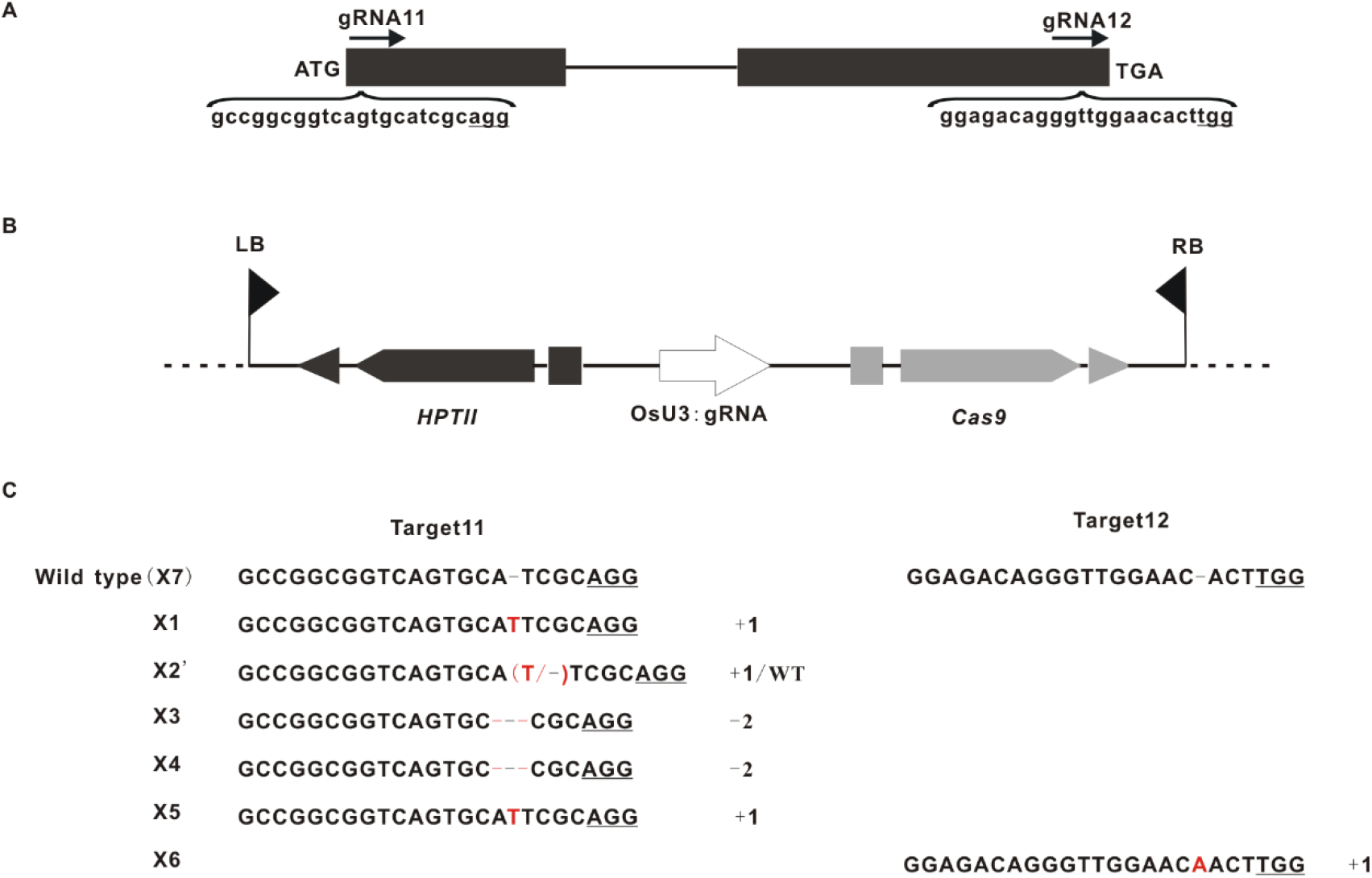
CRISPR/Cas9-induced *Pi-ta* gene modification in rice. (**A**) *Pi-ta* target loci. The gRNA11 and gRNA12 targeting sites were designed in the first and second exons of *Pi-ta*. The target sites are labeled in black lowercase letters. The protospacer adjacent motif (PAM) sequences are underlined. (**B**) The vector used to transform rice. gRNA11 and gRNA12 were assembled into the pGREB31 expression vector for Nipponbare transformation. (**C**) Nucleotide sequences at the target site in the 6 T_0_ mutant rice plants. The target site nucleotides are indicated as black capital letters and black dashes. The PAM site is underlined. The red dashes indicate the deleted nucleotides. The red capital letters indicate the inserted nucleotides. The numbers on the right indicate the number of nucleotides involved. “−” and “+” indicate the deletion and insertion of the indicated number of nucleotides, respectively.

In T_0_ transgenic plants, we screened six lines with targeted mutations (Figure 1C), five of which (X1, X3, X4, X5 and X6) showed homozygous mutations while only one (X2’) was heterozygous. A total of five mutations were achieved at Target 11, in contrary to the one mutation found at Target 12. These frameshift mutations would result in a loss-of-function of the *Pi-ta* gene. Of the six lines, X5 was eliminated from further analysis due to its high sterility.

Mutants of T_0_ were planted and 253 T_0_ plants were screened to analyze their genetic transformation pattern. T-DNA free mutants were selected using Cas9 gene-specific primers (Cas9S/Cas9AS, Supplementary table 1). The results showed that 42 mutants were not amplified to the Cas9 vector sequence, and were thus termed as T-DNA free plants. Of the T-DNA free plants, the inheritance of mutants at the target sites of lines X1, X3, X4 and X6 were further checked by sequencing. In X2’, which had heterozygous mutations at the target site, homozygous individuals and non-mutated individuals were screened in the segregating T1 siblings, termed as X2 and X0, respectively. In order to create rigorously controlled isogenic lines with the same genetic background, X0 was treated as a control without mutations (*Pi-ta*^0^) as it experienced the same cultivation, transformation and regeneration procedures as the other five *Pi-ta* knockout mutant lines (collectively, *Pi-ta*^−^). Nipponbare was included as the wild-type in this study, termed X7 (*Pi-ta*^WT^) herein. All the lines were then selfed to the T3 generation for further analysis.

### Fitness decline in *Pi-ta* knockout mutants in the absence of pathogens

To test the fitness effect of the *Pi-ta* gene in the absence of infection, we measured the relative fitness of *Pi-ta*^−^/*Pi-ta*^0^/*Pi-ta*^WT^ isolines (representing knockout, non-mutated and wild-type backgrounds, respectively) in field and greenhouse experiments in the absence of AVR-*Pita*.

In the field experiment, in 20 out of 25 comparisons using five fitness proxies for five independent knockout mutants, *Pi-ta*^−^ isogenic lines demonstrated significantly lower performance than *Pi-ta*^WT^ (Figure 2A-D; Supplementary figure 1A; Supplementary table 2). In terms of dry weight, height, number of filled grains and filled grains yield, *Pi-ta*^−^ isogenic lines suffered up to 41%, 28%, 42% and 49% reductions, respectively, relative to the *Pi-ta*^WT^ line (Figure 2A-D; Supplementary table 2).

**Figure 2.**
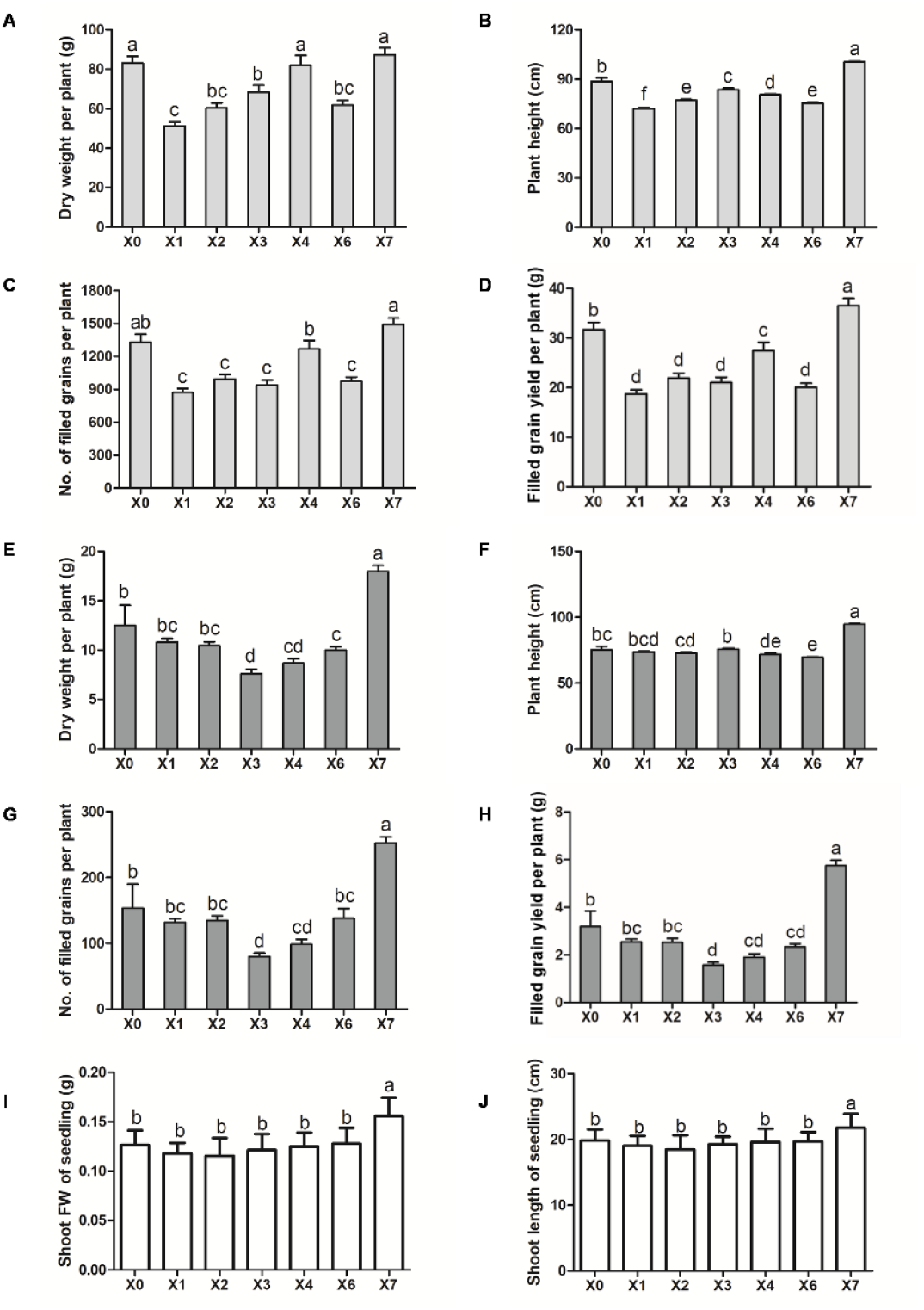
Significant fitness variations among isogenic lines in the absence of pathogen. X1, X2, X3, X4 and X6 are independent knockout mutant lines of the *Pi-ta* susceptible allele; X7 is the wild type background line; X0 is the non-mutated line screened in the X2 siblings, treated as the control without mutations in *Pi-ta*. Lines with different independent mutations in the *Pi-ta* gene were tested in the field, greenhouse experiments and in sterile condition. Bars with different letters are significantly different. Statistical differences among the agronomic traits were detected by Duncan’s multiple range test (*P* < 0.05). (**A-D**) Field fitness results. (**A**) Dry weight. (**B**) Height. (**C**) Number of filled grains. (**D**) Filled grain yield. (**E-H**) Greenhouse fitness results. (**E**) Dry weight. (**F**) Height. (**G**) Number of filled grains. (**H**) Filled grains yield. (**I-J**) Sterile condition fitness results. (**I**) Shoot fresh weight of seedling. (**J**) Shoot length of seedling. Values are means ± standard error (± SE).

In 19 of 25 comparisons using five fitness proxies for five independent knockout mutants, *Pi-ta*^−^ isogenic lines demonstrated significantly lower performance than *Pi-ta*^0^ (Figure 2A-D; Supplementary figure 1A; Supplementary table 2). In terms of dry weight, height, number of filled grains and filled grains yield, *Pi-ta*^−^ isogenic lines suffered up to 38%, 19%, 35% and 41% reductions, respectively, relative to the *Pi-ta*^WT^ line (Figure 2A-D; Supplementary table 2).

Taken together, these results demonstrate drastic fitness decline in *Pi-ta* knockout mutants in the absence of pathogens, which suggests that normal functioning of the *Pi-ta* susceptible allele in Nipponbare is beneficial in the absence of known *Pi-ta*-mediated pathogens carrying AVR-*Pita*.

To verify the robustness of our results, this fitness experiment was repeated in a greenhouse that was known to be free of *Pi-ta*-recognized pathogens. Similar to the field experiment, *Pi-ta*^−^ isogenic lines exhibited significantly lower performance in terms of dry weight, height, number of filled grains and filled grains yield relative to both *Pi-ta*^0^ and *Pi-ta*^WT^ isolines (Figure 2E-H; Supplementary figure 1B; Supplementary table 3). Thus, our greenhouse results recapitulated the results observed in the field.

One possibility to explain the observed fitness decline of all *Pi-ta* knockout mutants in the absence of *Magnaporthe oryzae* (AVR-*Pita*) is, that the presence of a different and undetected pathogen recognized by the allele of *Pi-ta* in *Pi-ta*^0^ or *Pi-ta*^WT^ isolines in the field may have provided a benefit to isolines carrying the normally functioning *Pi-ta* allele.

As a final confirmation that the observed fitness decline was not due to an interaction with an unknown pathogen, we grew our isogenic lines in sterile conditions on agar (Supplementary figure 2). Again, *Pi-ta*^0^ and *Pi-ta*^WT^ isolines had a higher weight and height than *Pi-ta*^−^ plants at 14 days (Figure 2I-J; Supplementary table 4). This result excluded the possibility that the presence of *Pi-ta* carried a fitness benefit because of the recognition of pathogens.

Collectively, these results demonstrate a beneficial function of the susceptible allele of *Pi-ta* in the absence of pathogens. Additionally, the results did not reveal significant variations in fitness between lines with different targeted mutation sites.

### *Pi-ta*-associated changes in defense response gene expression in the absence of pathogen

Expression levels can alter the penetrance of phenotypes (Raj, et al. 2010), and alternation of the expression of *Pi-ta* can lead to non-specific activation of the hypersensitive response (HR) or even lethality, if expression levels are too high (Wang, et al. 2019). To elucidate the molecular components involved in fitness decline in *Pi-ta* knockout mutants, we first evaluated the expression of the *Pi-ta* gene by qRT-PCR in the *Pi-ta*^−^ (X1, X2, X3, X4 and X6), *Pi-ta*^WT^ (X7), and *Pi-ta*^0^ (X0) lines in the absence of relevant pathogens (Figure 3A). In many cases, *Pi-ta* expression was 1.2- to 2.0-fold higher in these isolines than in the wild-type line, with the exception of X6. Expression of the *Pi-ta* gene was significantly lower in the X6 line compared with wild-type plants, suggesting considerable variation in the expression of the *Pi-ta* gene existed across targeted mutation sites. Replicated fitness decline induced by alternation in expression of the *Pi-ta* gene, whether higher or lower expression, suggested that this process was likely initiated at the post-transcriptional level.

**Figure 3.**
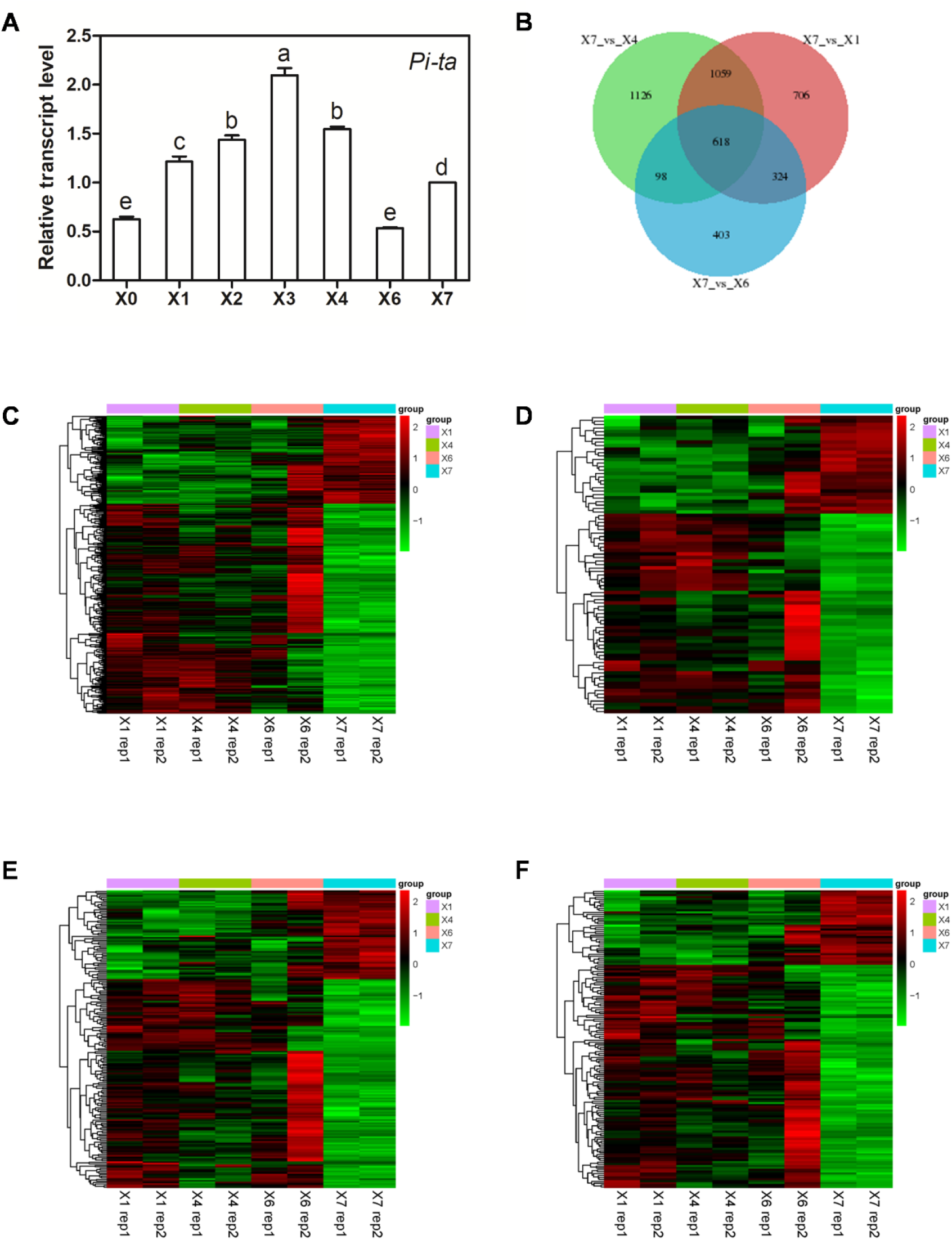
*Pi-ta* knockout mutant lines differentially express stimulus response, abiotic stimulus response, stress response and signal transduction genes relative to the wild type background. X1, X2, X3, X4 and X6 are independent knockout mutant lines of the *Pi-ta* susceptible allele; X7 is the wild type background line; X0 is the non-mutated line screened in the X2 siblings, treated as the control without mutations in *Pi-ta*. (**A**) Real-time PCR analysis of *Pi-ta* expression in leaves of 2-month-old plants grown in the greenhouse under standard conditions. Expression of *Pi-ta* in each line has been shown relative to that of *Pi-ta* in X7. Data are means ± SE. Bars with different letters are significantly different. Statistical differences among the relative transcriptional level were detected by Duncan’s multiple range test (*P* < 0.05). (**B**) The overlap of differentially expressed genes for three contrasts of lines with a *Pi-ta* allele and the knockout. (**C-F**) Heatmaps and dendrograms of gene sets as described below. Genes are in rows, and biological replicates are in columns, with both dendrograms grouped by similarity of expression in the gene set displayed. (**C**) Differentially expressed genes with GO annotations related to stimulus response. (**D**) Differentially expressed genes with GO annotations of response to abiotic stimulus. (**E**) Differentially expressed genes with GO annotations of response or stress. (**F**) Differentially expressed genes with GO annotations related to signal transduction.

The fitness data displayed two patterns that we further explored with transcriptome data. First, we observed that *Pi-ta* knockout lines had lower fitness than the *Pi-ta*^WT^ line. Second, *Pi-ta* knockout lines had lower fitness than the *Pi-ta*^0^ line. We first determined and compared the expression profile of three *Pi-ta*^−^ (X1, X4 and X6) lines and one *Pi-ta*^WT^ (X7) line, representing loss of function and normal function of the susceptible allele of *Pi-ta*, respectively. *Pi-ta*^−^ lines had 361 genes that were co-upregulated and 257 genes that were co-downregulated relative to the *Pi-ta*^WT^ line (Figure 3B). These genes were enriched for gene ontology (GO) annotations of response to stimulus, abiotic stimulus, stress, and signal transduction (Figure 3C, D, E, F and Supplementary table 5; *P* = 1.60 × 10 ^−6^, 4.00 × 10 ^−6^, 5.1 × 10 ^−4^, 2.8 × 10 ^−3^). We found substantial overlap in genes differentially expressed in independent contrasts of a single *Pi-ta*^−^ line and the *Pi-ta*^WT^ line (Figure 3B); genes identified in each single-line comparison were consistently enriched for the same GO annotations (Figure 3D; Supplementary table 6-9). Thus, loss of function of the susceptible allele from Nipponbare altered expression of genes that were involved in response to stress and signal transduction.

We explored the second of these two patterns by comparing the expression profiles of two background rigorously controlled isolines, X2 and X0, where only one nucleotide was inserted in the coding region of X2. The X2 line had 50 genes that were upregulated and 50 genes that were downregulated relative to the X0 line (Figure 4A). Genes upregulated in X2 plants were enriched for GO annotations involved in the integrin-mediated signaling pathway and mRNA cleavage (Supplementary table 9; *P* = 3.6 × 10 ^−3^, 1.76 × 10 ^−2^), whereas genes downregulated in X2 plants were enriched for inositol phosphate-mediated signaling, virus response, and hormone biosynthetic process annotations (Supplementary table 9; *P* = 3.6 × 10 ^−3^, 1.41 × 10 ^−2^, 2.3 × 10 ^−2^). In contrast to the above comparison between *Pi-ta*^−^ and *Pi-ta*^WT^, the X2 line differentially expressed an additional, unique set of genes involved in the hormone biosynthetic and mRNA cleavage processes in addition to signal transduction. Plant hormones, including jasmonic acid (JA), salicylic acid (SA) and ethylene (E), are all key regulators of defense responses (Liu, et al. 2014). The enrichment of hormone biosynthetic genes in the X2 line prompted us to examine JA, SA and 1-aminocyclopropane-1-carboxylic acid (ACC) as the precursor of E levels in both the X2 and X0 lines. As shown in figure 4B-D, JA, SA and ACC concentrations were significantly higher in X2 plants than in X0 plants (with fold changes ranging from 1.43 to 1.72, *P* < 0.05 by ANOVA). These data indicate that loss of function of the *Pi-ta* gene induced by one nucleotide insertion in the coding region caused major alterations in the endogenous levels of JA, SA and ACC.

**Figure 4.**
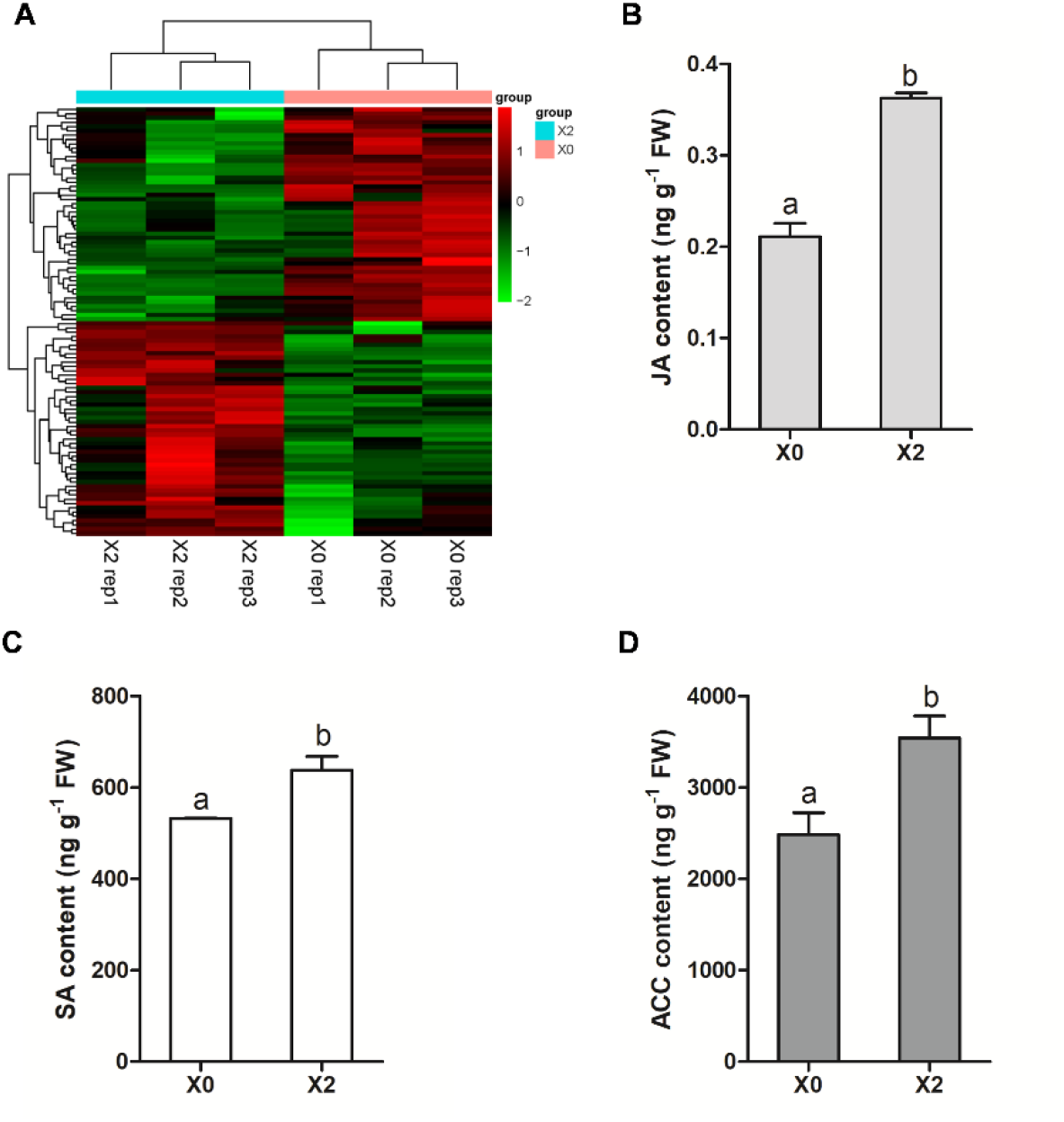
*Pi-ta* knockout mutant line X2 differentially express signal transduction, mRNA cleavage and hormone biosynthesis genes relative to its sibling line (X0) without mutation in *Pi-ta*. (**A**) Heatmaps and dendrograms of differentially expressed gene sets. Genes are in rows, and biological replicates are in columns, with both dendrograms grouped by similarity of expression in the gene set displayed. (**B-D**) Effect of *Pi-ta* knockout on accumulation of JA, SA and ACC. Three replicates of the X0 and X2 lines were grown in the greenhouse and harvested on days 60 of growth for determination of JA (**B**), SA (**C**) and ACC (**D**). Data are means ± SE. Bars with different letters are significantly different. Statistical differences among the relative transcriptional level were detected by Duncan’s multiple range test (*P* < 0.05).

## Discussion

### Drastic fitness decline in *Pi-ta* knockout mutants might be induced by autoimmunity

Gene expression can account for altered penetrance (Raj, et al. 2010), and alternation in the expression of the disease *R* gene often leads to non-specific activation of HR, or even lethality, if expression levels are too high (Wang, et al. 2019). In this study, the knockout mutants of the *Pi-ta* susceptible allele exhibited a significant decline in plant dry weight, height, number of filled grains and filled grain yield relative to both the wild type and non-mutated individuals. *Pi-ta* knockout in the japonica cultivar Wuyungeng24 also led to deleterious phenotypic effects, including shorter and weaker plants as well as frequent deaths (Wang, et al. 2019). This drastic fitness reduction suggests that the loss of function of the *Pi-ta* gene might induce overstimulation or mis-regulation of the immune system in rice.

Plant immunity is under tight control to avoid activation of defense responses in the absence of pathogens, as failure to do so can lead to autoimmunity that compromises plant growth and development. Many autoimmune mutants have been reported, most of which are associated with dwarfism and often spontaneous cell death (Rodriguez, et al. 2016). The rice *leaf11* gene, which encodes a protein with E3 ubiquitin ligase activity, induced chlorosis and spotted lesions on leaves due to constitutive activation of defense responses (Zeng, et al. 2004). The JA-governed defense response was strongly activated in *cea62* mutants, resulting in brown lesion spots over the entire leaf surface in *Arabidopsis* (Li, et al. 2012).

Plant immune receptors are generally maintained in inactive states and are activated only upon detection of pathogens. The benefit of carrying a functional allele of *Pi-ta* appears to result from its function as a negative regulator of the defense response, as loss of function of *Pi-ta* causes the upregulation of a number of genes involved in induced responses to stimulus and elevated hormones (Figure 3; Figure 4; Supplementary table 5-9). As the sensor confers the recognition of effectors, *Pi-ta* is essential for suppressing its helper (*Pi-42*)’s autoactivation in the absence of the pathogen, which could be deleterious to the host plant (Wang, et al. 2019).

Interestingly, morphological phenotypes resulting from plant autoimmunity often depend on the effects of genetic background. For example, the absence of elevated basal resistance and no other visible phenotypes were noticed in *srfr* (suppressor of *rps4*-RLD) mutants in the RLD ecotype, compared to that in the Col-0 ecotype (Kwon, et al. 2004). Moreover, no phenotypic changes have been observed in *BON1* (negative regulator of *SNC1*) mutants in the Ws ecotype compared to the dwarf phenotype in mutants of the Col-0 ecotype (Yang and Hua 2004). In this study, even though the *Pi-ta* knockout mutants in Nipponbare showed a drastic fitness decline including shorter, lighter plants and reduced grain yield, is still different from the frequent death reported in another japonica cultivar, Wuyungeng24 (Wang, et al. 2019). The effects of genetic background on growth-defense tradeoffs also suggest that different plant genotypes fine-tune their basal *R* gene expression levels.

### Fitness benefit plays a vital role in the retention of the *Pi-ta* susceptible alleles

In plants, a large number of *R* genes that segregate as loci with alternative alleles conferring different resistance to pathogens have been maintained over a long evolutionary time (Michelmore and Meyers 1998). As such, there would be no reason for hosts to harbor susceptible alleles in view of their lack of contribution to resistance. Why should rice populations still support disease-susceptible individuals of the *Pi-ta* gene along with disease-resistant individuals? There are several possible explanations for the long-term maintenance of these susceptible alternatives.

First, rice germplasm without a *Pi-ta* homolog has not been identified, suggesting that the *Pi-ta* gene might play physiological roles in addition to resistance to *M. oryzae* (Jia, et al. 2016). Furthermore, the discovery of potential multiple *Pi-ta* products from alternative splicing and exon skipping raises the possibility that they may play a role in recognizing diverse pathogen signals (Costanzo and Jia 2009).

Our creation of an artificial knockout of the *Pi-ta* susceptible allele in Nipponbare revealed a fitness decline of up to 49% upon the loss of *Pi-ta*’s function. The benefit of carrying a functional allele of *Pi-ta* appears to result from its function as a negative regulator of the defense response. Serving as an off-switch to its hepler Pi-42 and downstream immune signaling, *Pi-ta* was shown to be involved in the fine-tuning of the disease resistant response, as done by the alternative alleles of *Rps2* (Macqueen, et al. 2016). The critical function of *Pi-ta* provides a clear explanation for why the susceptible alleles have not been eliminated, as they typically are for *Rpm1* and *Rps5*. This result may help explain why the *Pi-ta* locus does not harbor true susceptible deletion mutants too.

Alternatively, the highly diversified *R* genes are assumed to persist as reservoirs for functionally distinct pathogen recognition alleles, providing sources for generating novel specificities by mutation and/or intergenic recombination (Michelmore and Meyers 1998). Multiple avirulent AVR-*Pita* haplotypes, determining the specificity of *Pi-ta*-mediated resistance, have been identified in commercial rice fields (Jia, et al. 2016). To keep pace with the fast evolution of the pathogens, more resistant *Pi-ta* alleles are in urgent need. Spontaneous mutations or reshuffling of variations in current alleles enables the continuous generation of diverse, novel alleles. Actually, the only one resistant *Pi-ta* haplotype detected thus far was inferred to derive from an ancestral haplotype H2 in wild rice (Huang, et al. 2008). Up to 100 haplotypes of the *Pi-ta* gene discovered in *Oryza* species might serve as an enlarged reservoir of variations to fight against various pathogens.

Finally, different from the wild plants, in arable crops, yield, quality and agronomic properties, such as optimal height and heading time, are all normally considered more important than disease resistance (Brown 2002). The need to prioritize other traits over disease resistance may have an impact on plant breeding programs. If resistance to a disease is not an important commercial target, if the susceptible alleles bring out fitness benefits such as yield increase [as inferred from Wang et al. (2008)], and if breeders therefore select for it, the frequency of the susceptible alleles in breeders’ germplasm will increase as a result of artificial selection. Thus, in over 10,000 years’ domestication of cultivated rice, plant breeding can be regarded as an essential force for the retention of the susceptible *Pi-ta* alleles (Zhao 2010).

Our results stood in consistency to the fitness benefits of *Rps2* in *Arabidopsis* (Macqueen, et al. 2016). They both showed that the *R* gene alleles, including the susceptible one, contribute to the regulation of gene expression and thus present a pleiotropic effect, explaining their maintenance in the genome.

Experimental studies to evaluate the fitness effect of *R* genes, as described above, have been focusing on loci with very simple genetic architectures. *R* genes show a high degree of copy number variation within and across plant species, and many of them are organized in tandem arrays (Jacob, et al. 2013). How, then, do plants with a vast repertoire of *R* genes coordinate the fitness costs and benefits of the alternative alleles will surely be a rich seam worthy of further mining.

## Materials and Methods

### Plant material and growth

The background of transgenic plants was Nipponbare (*Oryza sativa* L. ssp. *Japonica*). All rice plants were grown in a greenhouse at our institute at 28-35°C or fields in our experimental station under normal growth conditions in Nanjing. The experimental station is specialized for genetically modified crop planting permitted by the Chinese Ministry of Agriculture.

### Vector construction

The pTGE1 vector was constructed according to a method described by Xie and Yang (2013). Briefly, the primers g11S/g11AS covering target site sequence 11 (Figure 1A; Supplementary table 1) were synthesized by Genscript (www.genscript.com.cn) and combined by annealing. The corresponding primers were cloned into the sgRNA expression cassette of pGREB31 (Addgene). The pTGE2 vector was constructed as described above based on target site sequence 12 (Figure 1B; Supplementary table 1).

### Rice transformation

The Cas9/sgRNA-expressing binary vectors (pTGE1 and pTGE2) were transformed into the *Agrobacterium tumefaciens* strain LBA4404 by electroporation. Agrobacterium-mediated transformation of embryogenic calli derived from ‘Nipponbare’ was performed according to procedures detailed in Hiei et al. (1994). Briefly, hygromycin-containing medium was used to select hygromycin-resistant calli, and then the hygromycin-resistant calli were transferred to regeneration medium for the regeneration of transgenic plants. After 2-3 months of cultivation, transgenic seedlings were transferred to the greenhouse until maturity.

### Identification of mutant transgenic plants

Genomic DNA was extracted from individual transgenic plants using SDS extraction (Dellaporta, et al. 1983). All transgenic hygromycin-resistant T_0_ plants were characterized by PCR using the Cas9-specific primers Cas9S/Cas9AS (Figure 1; Supplementary table 1). Subsequently, all PCR-positive plants were subjected to PCR using the gene-specific primer pairs, *Pi-ta*787/*Pi-ta*1900 and *Pi-ta*4678/*Pi-ta*5522 (Figure 1; Supplementary table 1), to amplify DNA fragments across the two target sites, respectively. The resulting PCR amplicons were then directly sequenced. The sequencing chromatograms with superimposed peaks of biallelic and heterozygous mutations were decoded by the Degenerate Sequence Decoding (DSD) method-based web tool DSDecode (http://dsdecode.scgene.com/) (Liu, et al. 2015).

The CRISPR-GE online tool (http://skl.scau.edu.cn/) was used to identify potential off-target sites for the two Cas9 sgRNAs from those edited plants. No sgRNAs had predicted off-target sites with fewer than 2-nt mismatches, and only one off-target site was predicted with 7-nt mismatches for sgRNA11, suggesting that the designed spacers were highly specific. For this one potential off-target site, PCR amplification followed by Sanger sequencing was used to check whether mutations were induced in all edited lines (Supplementary table 1). To further assess possible off-target effects, one (X2) of five edited lines was arbitrarily selected for whole genome sequencing and the mutations were identified using a pipeline described in Wang et al. (2019). The targeted mutation was confirmed in the sequenced plant, and no novel mutations were found in predicted off-target sites even when up to 8-nt mismatched were allowed, suggesting a very low off-targeting rate.

The presence of CRISPR/Cas9 DNA and marker genes in gene-edited plants may cause adverse effects, such as an increased risk of off-target changes, and may trigger regulation concerns when these plants are used in crop breeding (Chen, et al. 2019). Here, we tried to screen for T-DNA free individuals in the T1 generation. The T-DNA free lines were screened by amplifying the genomic DNA of T1 generation lines using Cas9 gene-specific primers (Cas9S/Cas9AS) (Supplementary table 1). The T-DNA free lines were selected for sequencing of the target regions. T-DNA-free lines with homozygous and no mutations were obtained for further agronomic trait characterization.

### Field and greenhouse fitness experiments

To reveal evolutionarily relevant fitness effects of resistance, fitness traits were assessed under natural growing conditions and in the absence of pathogens. Seedlings of each of seven *Pi-ta* lines were grown in the field under normal growth conditions in Nanjing. The experimental plot was partitioned into 20 blocks in a randomized block design, in which each block contained four replicates for each *Pi-ta* line. Each block consisted of four rows with seven plants per row, with one meter (m) between rows and 0.9 m spaces within rows. The field was hand weeded once and plants received no other protection from competition or pests. Roughly 12% of plants died, but these were evenly distributed among the lines.

The greenhouse fitness experiment was performed in 20 square buckets. Each bucket contained 4 replicates from each *Pi-ta* line in a randomized design. Plants were set out in four rows in each bucket, spaced by 0.3 m within rows and by 0.4 m between rows. Roughly 20% of plants died, but these were evenly distributed among the lines.

Five fitness proxies were measured in the field and the greenhouse fitness experiments: dry weight, height, effective tillers, number of filled grains and filled grain yield.

All seven *Pi-ta* lines were grown in sterile conditions. The sterile experiments included 10 seedlings per line. Seeds were surface sterilized by briefly vortexing in 75% ethanol and then soaked in 30% (v/v) sodium hypochlorite for 5 min and thoroughly washed. Seeds were germinated on ½MS media with agar in sterile beakers, and seven seeds from different lines were evenly distributed in one beaker in a randomized design. Two-week old seedlings were harvested to measure plant height, fresh weight and dry weight.

### Sampling for pathogen presence

A key element to accurately measure the costs of resistance is to confirm that no pathogens that would have interfered with the experiment were present at the field site.

Ninety-six plant samples representing all seven lines were surface sterilized and plated on PAD medium on days 30 and 60 of growth during the field and greenhouse experiments. Out of hundreds of colonies, 42 potentially distinct fungus types were identified by general appearance. Each type was sequenced for ITS (Khang, et al. 2008) and compared to published sequences in GenBank. None of the fungus isolates were *Magnaporthe oryzae* according to their ITS identity. Finally, we used PCR to screen for the presence of AVR-*Pita*. We screened with four combinations of primer pairs (Supplementary table 1) based on the published sequence, but AVR-*Pita* was not detected in any of our colonies.

### Quantification of salicylic acid (SA), jasmonic acid (JA) and 1-aminocyclopropane-1-carboxylic acid (ACC)

Three replicates of X0 and X2 lines were grown in the greenhouse and harvested on day 60 of growth.

Jasmonic acid was extracted and quantified as described (Schweizer, et al. 1997). Briefly, 0.5-1.0 g of fresh leaves were used for JA purification and GC/MS analysis (Finnigan Trace GC–MS, USA). 9,10-dihydro-JA was added as an internal standard.

SA was extracted from 1.0 g of fresh leaves and quantified with an HPLC system equipped with fluorescence detection (LC-2010AHT, Shimadzu) as described (Metwally, et al. 2003). Authentic SA was used for calibration.

1-aminocyclopropane-1-carboxylic acid (ACC) was extracted and quantified as described (Concepcion, et al. 1979). One gram of fresh leaves was used for ACC purification and HPLC-MS analysis. Authentic ACC was used for calibration.

### Whole transcriptome profiling

Two or three biological replicates (10 plants each) of wild-type and *Pi-ta* mutant plants were sampled for RNA sequencing. Total RNA was extracted using Trizol Reagent (Life Technologies). The qualified RNA samples were then used for library construction following the specifications of the TruSeq RNA Sample Preparation v2 Guide (Illumina), and RNA sequencing was conducted on an Illumina Hiseq 2500 at Shanghai Personal Biotechnology Co., Ltd. (Shanghai, China). We used SeqPrep to strip adaptors and/or merge paired reads that overlapped into single reads and used Sickle to remove low-quality reads. The clean data were then mapped to the reference genome of rice using HISAT2 v2.1.0. FPKM (fragments per kilobase per million mapped reads) was then calculated to estimate the expression levels of genes. DESeq2 v1.6.3 was used to analyze the differential gene expression between two samples, and genes with q < 0.05 and |log2_ratio| > 1 were identified as differentially expressed genes. GO (Gene Ontology; http://geneontology.org/) enrichment of the DEGs was implemented by hypergeometric tests, in which the p-value was calculated and adjusted as the q-value. GO terms with q < 0.05 were considered significantly enriched.

### Quantitative Reverse Transcription Polymerase Chain Reaction (qRT-PCR) Analysis

Leaves were sampled from 2-month-old plants grown in the greenhouse under standard conditions. Total RNA isolation, first-strand cDNA synthesis and qRT-PCR analysis were performed as described previously (Sharma, et al. 2013). Three PCR replicates were run for each line. Experiments were conducted in three biological replicates for each sample and three technical replicates were analyzed for each biological replicate. The ΔΔC_T_ calculation method was used to calculate the relative expression level of each gene. For normalizing the relative mRNA level of individual gene in various RNA samples, EF-257 was used as an internal control gene (Yang, et al. 2018). The list of primer sequences used for qRT-PCR analysis is provided in Supplementary table 1.

### Statistical analysis

Data analysis was performed in Microsoft Excel and is presented as the mean ±standard error (SE) of biological replicates. Statistical analysis was performed using One-way-Analysis-of Variance (ANOVA); this was followed by Duncan’s test using SPSS (version 19.0) for data statistics. Different letters are used (*P* < 0.05) to present statistically significant differences and similar letters are considered as statistically non-significant.

## Acknowledgements

We thank Dr. Dacheng Tian for donation of rice material. This work was supported by grants from the National Natural Science Foundation of China to X.Q.S. (Grant no. 31470448), Jiangsu Key Laboratory of Plant Resources Research and Utilization grant to X.Q.S (JSPKLB201921), Changshu Agricultural Production and Public Service Project and Doctoral Research Foundation supported by the Institute of Botany, Jiangsu Province and Chinese Academy of Sciences to J.L. (JSPKLB201906).

