## Supplementary figures and tables for "Fitness benefit plays a vital role in the retention of the *Pi-ta* susceptible alleles"

**Supplementary Figure 1. Effective tillers variations among isogenic lines in the absence of pathogen.** X1, X2, X3, X4 and X6 are independent knockout mutant lines of the *Pi-ta* susceptible allele; X7 is the wild type background line; X0 is the non-mutated line screened in the X2 siblings, treated as the control without mutations in *Pi-ta*. Lines with different independent mutations in the *Pi-ta* gene were tested in both the field and the greenhouse experiments. Bars with different letters are significantly different. Statistical differences among the agronomic traits were detected by Duncan's multiple range test ($p < 0.05$). a, Field results. b, Greenhouse results.

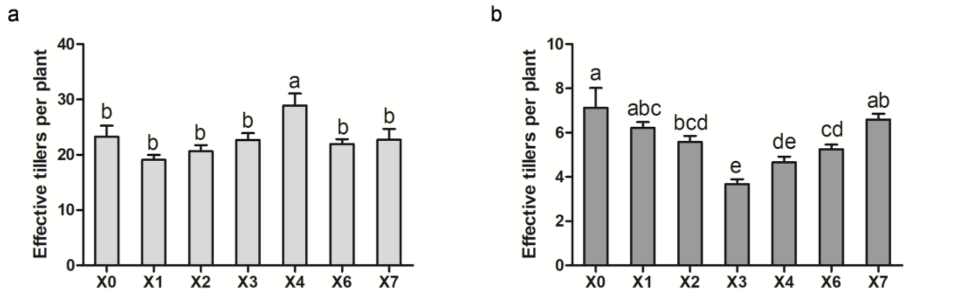

**Supplementary Figure 2. Seedling phenotype of isogenic lines at 14 days in sterile conditions.** X1, X2, X3, X4 and X6 are independent knockout mutant lines of the *Pi-ta* susceptible allele; X7 is the wild type background line; X0 is the non-mutated line screened in the X2 siblings, treated as the control without mutations in *Pi-ta*. Scale bars = 5 cm.

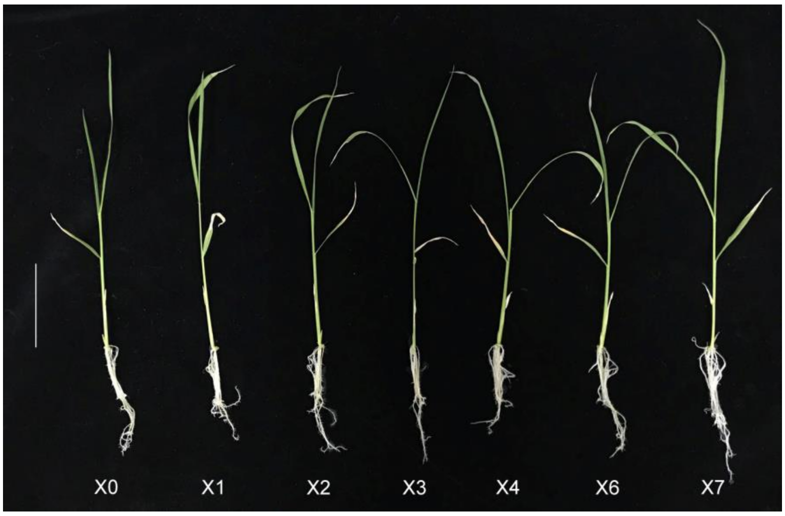

**Supplementary Table 1. List of PCR primer pairs used in this study.**

| Primer | Sequence | Purpose |
|---|---|---|
| **g11S** | 5'-ggcaGCCGGCGGTCAGTGCATCGC-3' | guide RNA11 synthesis |
| **g11AS** | 5'-aaacGCGATGCACTGACCGCCGGC-3' | |
| **g12S** | 5'-ggcaGGAGACAGGGTTGGAACACT-3' | guide RNA12 synthesis |
| **g12AS** | 5'-aaacAGTGTTCCAACCCTGTCTCC-3' | |
| **Cas9S** | 5'-TGGTGGAAGAGGATAAGAAGC-3’ | PCR analysis on the Cas9 gene |
| **Cas9AS** | 5’-TCAAACAGTGTCAGGGTCAGC-3’ | |
| ***Pita*787** | 5’-TGCCTACTTCTTTCCACATCA-3’ | Cloning and sequencing of target 11 |
| **Pita1900** | 5’-ATCCGAAGACTGATGCTGATGTT-3’ | |
| **Pita4678** | 5'-CAGGATGACCTTGACACTCTC-3' | Cloning and sequencing of target12 |
| **Pita5522** | 5'- GCATACACCTTTCGTTCTTTG-3' | |
| **11offS** | 5'-ACTGCGTCAAGTGGTCAAGC-3' | Cloning and sequencing of the |
| **11offAS** | 5'-CACACAACGGATTTCCAGGG-3' | potential offsite target |
| **ITS1** | 5'-TCCGTAGGTGAACCTGCGG-3' | Fungus identification |
| **ITS4** | 5'-TCCTCCGCTTATTGATATGC-3' | (White et al., 1990) |
| **AVR-Pita1-F** | 5'-AATCGACAGTTTTGTTTGCACAATC-3' | PCR analysis on the AVR-Pita gene |
| **AVR-Pita1-R** | 5'-TCCCTCCATTCCAACACTAACG-3' | |
| **AvrPita-F** | 5'-CAGGCATACATTGGAGAGCC-3' | PCR analysis on the AVR-Pita gene |
| **AvrPita-R** | 5'-CCCTCCATTCCAACACTAAC-3' | (IRRI) |
| **q_Os*Pi-ta* F3** | 5'-GAAGCCTGACATGACGAAGATC-3' | qRT-PCR analysis on the *Pi-ta* gene |
| **q_Os*Pi-ta* R3** | 5'-CAAACCACGGCTAACAATATCC-3' | (Yang et al.,2017) |

**Supplementary Table 2. Effect of *Pi-ta* functioning or loss-function on correlates of fitness in field experiments.**

| Lines | Dry Weight/g | Height/cm | Effective Tillers | Number of Filled Grains | Filled Grain Yield/g |
|---|---|---|---|---|---|
| **X0** | 83.15±3.45a | 88.76±2.11b | 23.29±1.94b | 1331.47±72.06ab | 31.70±1.42b |
| **X1** | 51.24±1.94c | 72.13±0.48f | 19.09±0.90b | 870.24±37.27c | 18.71±0.84d |
| **X2** | 60.38±2.48bc | 77.28±0.44e | 20.66±1.08b | 993.50±42.34c | 21.95±0.92d |
| **X3** | 68.44±3.42b | 83.73±0.87c | 22.68±1.24b | 939.96±44.17c | 21.03±1.04d |
| **X4** | 81.86±5.22a | 80.62±0.49d | 28.93±2.17a | 1269.14±77.50b | 27.44±1.69c |
| **X6** | 61.85±2.34bc | 75.41±0.53e | 21.94±0.88b | 973.88±37.93c | 20.05±0.83d |
| **X7** | 87.39±3.48a | 100.63±0.44a | 22.72±1.96b | 1496.38±58.32a | 36.56±1.43a |

Results are means ± s.e. Values with different letters are statistically significant and those with the same letters are not significantly different ($P < 0.05$).

**Supplementary Table 3. Effect of *Pi-ta* functioning or loss-function on correlates of fitness in greenhouse experiments.**

| Lines | Dry Weight/g | Height/cm | Effective Tillers | Number of Filled Grains | Filled Grain Yield/g |
|---|---|---|---|---|---|
| **X0** | 12.51±2.03b | 75.13±3.00bc | 7.13±0.90a | 153.38±36.61b | 3.19±0.66b |
| **X1** | 10.80±0.39bc | 73.59±0.55bcd | 6.22±0.26abc | 131.92±5.94bc | 2.54±0.12bc |
| **X2** | 10.45±0.39bc | 72.63±0.87cd | 5.58±0.26abc | 134.83±6.92bc | 2.54±0.16bc |
| **X3** | 7.62±0.41d | 75.84±0.72b | 3.68±0.21e | 80.06±5.34d | 1.58±0.11d |
| **X4** | 8.68±0.47cd | 71.68±1.13de | 4.65±0.27de | 98.55±7.56cd | 1.89±0.15cd |
| **X6** | 9.99±0.38c | 69.54±0.44e | 5.25±0.21cd | 138.42±14.37bc | 2.34±0.13cd |
| **X7** | 17.98±0.62a | 94.76±0.53a | 6.60±0.25ab | 252.30±9.41a | 5.75±0.22a |

Results are means ± s.e. Values with different letters are statistically significant and those with the same letters are not significantly different ($P < 0.05$).

**Supplementary Table 4. Effect of *Pi-ta* functioning or loss-function on correlates of fitness at 14 days in sterile conditions.**

| Lines | Shoot fresh weight of seedling /g | Shoot length of seedling /cm |
|---|---|---|
| **X0** | 0.1264±0.0047b | 19.86±0.53b |
| **X1** | 0.1178±0.0034b | 19.04±0.48b |
| **X2** | 0.1156±0.0060b | 18.46±0.74b |
| **X3** | 0.1214±0.0054b | 19.27±0.39b |
| **X4** | 0.1250±0.0045b | 19.61±1.13b |
| **X6** | 0.1277±0.0054b | 19.69±0.47b |
| **X7** | 0.1556±0.0060a | 21.82±0.66a |

Results are means ± s.e. Values with different letters are statistically significant and those with the same letters are not significantly different ($P < 0.05$).

**Supplementary Table 5. Gene ontology enrichment of the 618 genes differentially expressed between the *Pi-ta*$^{-}$ (X1, X4, X6) and *Pi-ta*$^{WT}$ (X7) lines.** *Pi-ta*$^{-}$ and *Pi-ta*$^{WT}$ lines are the *Pi-ta* knockout lines and the wild type line respectively. GO annotations are from experimentally verified datasets and differentially expressed genes were compared to the entire annotated *Oryza sativa* genome. Percentage is the number of genes enriched in one GO pathway divided the total number of genes differentially expressed between the *Pi-ta*$^{-}$ and *Pi-ta*$^{WT}$ lines.

| GO annotation | # DE in *Pi-ta*$^{-}$ vs *Pi-ta*$^{WT}$ | Percentage | # in Genome | *P* value |
|---|---|---|---|---|
| **response to stimulus** | 359 | 0.580906149 | 3356 | 1.60E-06 |
| **response to abiotic stimulus** | 85 | 0.137540453 | 604 | 4.00E-06 |
| **positive regulation of chromosome** | 6 | 0.009708738 | 8 | 9.80E-06 |
| **asymmetric cell division** | 6 | 0.009708738 | 8 | 9.80E-06 |
| **DNA geometric change** | 19 | 0.030744337 | 77 | 2.10E-05 |
| **DNA duplex unwinding** | 19 | 0.030744337 | 77 | 2.10E-05 |
| **positive regulation of histone modification** | 5 | 0.008090615 | 6 | 2.60E-05 |
| **positive regulation of chromatin organization** | 5 | 0.008090615 | 6 | 2.60E-05 |
| **microtubule-based movement** | 21 | 0.033980583 | 93 | 3.30E-05 |
| **movement of cell or subcellular component** | 21 | 0.033980583 | 98 | 7.50E-05 |
| **regulation of fatty acid biosynthetic process** | 4 | 0.006472492 | 5 | 0.00026 |
| **negative regulation of protein acetylation** | 4 | 0.006472492 | 5 | 0.00026 |
| **negative regulation of peptidyl-lysine** | 4 | 0.006472492 | 5 | 0.00026 |
| **negative regulation of histone acetylation** | 4 | 0.006472492 | 5 | 0.00026 |
| **DNA conformation change** | 26 | 0.042071197 | 148 | 0.00036 |
| **response to stress** | 191 | 0.309061489 | 1773 | 0.00051 |
| **positive regulation of histone methylation** | 3 | 0.004854369 | 3 | 0.00064 |
| **regulation of protein acetylation** | 4 | 0.006472492 | 6 | 0.00072 |
| **regulation of peptidyl-lysine acetylation** | 4 | 0.006472492 | 6 | 0.00072 |
| **regulation of histone acetylation** | 4 | 0.006472492 | 6 | 0.00072 |
| **regulation of fatty acid metabolic process** | 4 | 0.006472492 | 6 | 0.00072 |
| **cellular response to stimulus** | 235 | 0.3802589 | 2258 | 0.00082 |
| **response to salicylic acid** | 8 | 0.012944984 | 25 | 0.00085 |
| **response to water** | 19 | 0.030744337 | 101 | 0.00092 |
| **cell cycle** | 52 | 0.084142395 | 395 | 0.00139 |
| **negative regulation of histone modification** | 4 | 0.006472492 | 7 | 0.00155 |
| **phosphorus metabolic process** | 279 | 0.451456311 | 2768 | 0.00178 |
| **DNA metabolic process** | 75 | 0.121359223 | 625 | 0.00204 |
| **peptidyl-histidine phosphorylation** | 5 | 0.008090615 | 12 | 0.00223 |

| | | | | |
|---|---|---|---|---|
| **cold acclimation** | 5 | 0.008090615 | 12 | 0.00223 |
| **cell communication** | 172 | 0.278317152 | 1627 | 0.00226 |
| **phosphorylation** | 211 | 0.341423948 | 2045 | 0.00237 |
| **DNA replication initiation** | 7 | 0.011326861 | 23 | 0.00248 |
| **response to acid chemical** | 42 | 0.067961165 | 312 | 0.00252 |
| **chromosome organization** | 60 | 0.097087379 | 483 | 0.00253 |
| **lipid transport** | 17 | 0.027508091 | 94 | 0.00259 |
| **DNA replication** | 21 | 0.033980583 | 127 | 0.00275 |
| **phosphate-containing compound metabolic** | 273 | 0.441747573 | 2724 | 0.00278 |
| **signal transduction** | 156 | 0.252427184 | 1466 | 0.0028 |
| **response to external stimulus** | 40 | 0.064724919 | 296 | 0.00294 |
| **signaling** | 156 | 0.252427184 | 1468 | 0.00295 |
| **xyloglucan metabolic process** | 11 | 0.017799353 | 50 | 0.00302 |
| **response to salt stress** | 21 | 0.033980583 | 129 | 0.00334 |
| **gravitropism** | 5 | 0.008090615 | 13 | 0.00337 |
| **microtubule-based process** | 26 | 0.042071197 | 174 | 0.004 |
| **response to water deprivation** | 17 | 0.027508091 | 98 | 0.00405 |
| **response to oxygen-containing compound** | 54 | 0.087378641 | 435 | 0.00409 |
| **protein kinase C-activating G-protein coupled** | 4 | 0.006472492 | 9 | 0.00486 |
| **regulation of histone modification** | 5 | 0.008090615 | 14 | 0.00488 |
| **DNA repair** | 42 | 0.067961165 | 325 | 0.00528 |
| **regulation of gene expression by genetic** | 3 | 0.004854369 | 5 | 0.00559 |
| **radial pattern formation** | 3 | 0.004854369 | 5 | 0.00559 |
| **protein phosphorylation** | 168 | 0.27184466 | 1622 | 0.00575 |
| **cellular response to stress** | 70 | 0.113268608 | 602 | 0.00602 |
| **DNA recombination** | 24 | 0.038834951 | 163 | 0.00666 |
| **cellular response to hormone stimulus** | 41 | 0.066343042 | 320 | 0.00671 |
| **response to gravity** | 5 | 0.008090615 | 15 | 0.0068 |
| **regulation of lipid biosynthetic process** | 5 | 0.008090615 | 15 | 0.0068 |
| **regulation of cellular ketone metabolic process** | 5 | 0.008090615 | 15 | 0.0068 |
| **peptidyl-histidine modification** | 5 | 0.008090615 | 15 | 0.0068 |
| **response to lipid** | 31 | 0.050161812 | 227 | 0.00701 |
| **regulation of lipid metabolic process** | 6 | 0.009708738 | 21 | 0.0071 |
| **phospholipid dephosphorylation** | 6 | 0.009708738 | 21 | 0.0071 |
| **phosphatidylinositol dephosphorylation** | 6 | 0.009708738 | 21 | 0.0071 |
| **positive regulation of protein modification** | 14 | 0.022653722 | 79 | 0.00714 |
| **DNA-dependent DNA replication** | 14 | 0.022653722 | 79 | 0.00714 |
| **response to temperature stimulus** | 25 | 0.040453074 | 173 | 0.00723 |
| **transposition, RNA-mediated** | 2 | 0.003236246 | 2 | 0.00742 |
| **secondary growth** | 2 | 0.003236246 | 2 | 0.00742 |

| | | | | |
|---|---|---|---|---|
| **response to herbivore** | 2 | 0.003236246 | 2 | 0.00742 |
| **regulation of tetrapyrrole catabolic process** | 2 | 0.003236246 | 2 | 0.00742 |
| **regulation of jasmonic acid biosynthetic** | 2 | 0.003236246 | 2 | 0.00742 |
| **regulation of iron ion transport** | 2 | 0.003236246 | 2 | 0.00742 |
| **regulation of histone H4 acetylation** | 2 | 0.003236246 | 2 | 0.00742 |
| **regulation of chlorophyll catabolic process** | 2 | 0.003236246 | 2 | 0.00742 |
| **pyrimidine dimer repair** | 2 | 0.003236246 | 2 | 0.00742 |
| **positive regulation of histone H3-K9** | 2 | 0.003236246 | 2 | 0.00742 |
| **pigmentation** | 2 | 0.003236246 | 2 | 0.00742 |
| **pigment accumulation in tissues in response to** | 2 | 0.003236246 | 2 | 0.00742 |
| **pigment accumulation in tissues** | 2 | 0.003236246 | 2 | 0.00742 |
| **pigment accumulation in response to UV light** | 2 | 0.003236246 | 2 | 0.00742 |
| **pigment accumulation** | 2 | 0.003236246 | 2 | 0.00742 |
| **negative regulation of iron ion transport** | 2 | 0.003236246 | 2 | 0.00742 |
| **negative regulation of histone H4 acetylation** | 2 | 0.003236246 | 2 | 0.00742 |
| **negative regulation of fatty acid metabolic** | 2 | 0.003236246 | 2 | 0.00742 |
| **negative regulation of fatty acid biosynthetic** | 2 | 0.003236246 | 2 | 0.00742 |
| **maintenance of protein location in nucleus** | 2 | 0.003236246 | 2 | 0.00742 |
| **maintenance of chromatin silencing** | 2 | 0.003236246 | 2 | 0.00742 |
| **lateral growth** | 2 | 0.003236246 | 2 | 0.00742 |
| **anthocyanin accumulation in tissues in** | 2 | 0.003236246 | 2 | 0.00742 |
| **cell wall biogenesis** | 19 | 0.030744337 | 121 | 0.00753 |
| **response to osmotic stress** | 22 | 0.035598706 | 147 | 0.00754 |
| **negative regulation of chromatin organization** | 4 | 0.006472492 | 10 | 0.00755 |
| **response to alcohol** | 23 | 0.037216828 | 156 | 0.00766 |
| **hormone-mediated signaling pathway** | 40 | 0.064724919 | 313 | 0.00767 |
| **cell division** | 24 | 0.038834951 | 165 | 0.00774 |
| **nucleobase-containing compound** | 231 | 0.373786408 | 2319 | 0.00803 |
| **cellular response to endogenous stimulus** | 41 | 0.066343042 | 324 | 0.00826 |
| **cellular response to DNA damage stimulus** | 43 | 0.069579288 | 344 | 0.00858 |
| **lipid localization** | 17 | 0.027508091 | 106 | 0.00902 |
| **pattern specification process** | 10 | 0.01618123 | 50 | 0.00928 |
| **regulation of cellular process** | 376 | 0.608414239 | 3938 | 0.00998 |
| **post-embryonic plant morphogenesis** | 12 | 0.019417476 | 66 | 0.01002 |
| **auxin-activated signaling pathway** | 16 | 0.025889968 | 99 | 0.01032 |
| **negative regulation of lipid metabolic process** | 3 | 0.004854369 | 6 | 0.01047 |
| **negative regulation of lipid biosynthetic** | 3 | 0.004854369 | 6 | 0.01047 |
| **genetic imprinting** | 3 | 0.004854369 | 6 | 0.01047 |
| **response to abscisic acid** | 22 | 0.035598706 | 152 | 0.01105 |
| **response to chitin** | 4 | 0.006472492 | 11 | 0.01107 |

| | | | | |
|---|---|---|---|---|
| **shoot system development** | 34 | 0.055016181 | 263 | 0.01118 |
| **regionalization** | 9 | 0.014563107 | 44 | 0.01147 |
| **G-protein coupled receptor signaling pathway** | 7 | 0.011326861 | 30 | 0.01212 |
| **cellular response to auxin stimulus** | 16 | 0.025889968 | 101 | 0.01242 |
| **regulation of nucleobase-containing** | 197 | 0.318770227 | 1973 | 0.01299 |
| **response to hormone** | 60 | 0.097087379 | 522 | 0.01309 |
| **aromatic compound biosynthetic process** | 248 | 0.401294498 | 2533 | 0.0133 |
| **positive regulation of organelle organization** | 6 | 0.009708738 | 24 | 0.0141 |
| **regulation of biological process** | 402 | 0.650485437 | 4255 | 0.01435 |
| **regulation of RNA biosynthetic process** | 190 | 0.307443366 | 1902 | 0.01437 |
| **regulation of nucleic acid-templated** | 190 | 0.307443366 | 1902 | 0.01437 |
| **response to organic substance** | 72 | 0.116504854 | 647 | 0.01448 |
| **intracellular signal transduction** | 47 | 0.07605178 | 395 | 0.01474 |
| **regulation of RNA metabolic process** | 193 | 0.312297735 | 1937 | 0.01504 |
| **response to endogenous stimulus** | 60 | 0.097087379 | 526 | 0.01517 |
| **mRNA transcription** | 4 | 0.006472492 | 12 | 0.01549 |
| **coenzyme A metabolic process** | 4 | 0.006472492 | 12 | 0.01549 |
| **regulation of transcription, DNA-templated** | 188 | 0.30420712 | 1889 | 0.01703 |
| **chaperone mediated protein folding requiring** | 3 | 0.004854369 | 7 | 0.01716 |
| **organic cyclic compound biosynthetic process** | 256 | 0.414239482 | 2640 | 0.01822 |
| **response to chemical** | 120 | 0.194174757 | 1161 | 0.01935 |
| **signal transduction by protein** | 12 | 0.019417476 | 72 | 0.01945 |
| **biological regulation** | 448 | 0.724919094 | 4798 | 0.0196 |
| **tropism** | 5 | 0.008090615 | 19 | 0.01973 |
| **RNA biosynthetic process** | 201 | 0.325242718 | 2039 | 0.01973 |
| **cutin biosynthetic process** | 5 | 0.008090615 | 19 | 0.01973 |
| **multicellular organism development** | 85 | 0.137540453 | 792 | 0.01983 |
| **negative regulation of cell cycle** | 9 | 0.014563107 | 48 | 0.01992 |
| **guard cell differentiation** | 4 | 0.006472492 | 13 | 0.02087 |
| **regulation of jasmonic acid metabolic process** | 2 | 0.003236246 | 3 | 0.02099 |
| **regulation of cell cycle arrest** | 2 | 0.003236246 | 3 | 0.02099 |
| **positive regulation of protein acetylation** | 2 | 0.003236246 | 3 | 0.02099 |
| **positive regulation of peptidyl-lysine** | 2 | 0.003236246 | 3 | 0.02099 |
| **positive regulation of MAPK cascade** | 2 | 0.003236246 | 3 | 0.02099 |
| **positive regulation of MAP kinase activity** | 2 | 0.003236246 | 3 | 0.02099 |
| **positive regulation of histone acetylation** | 2 | 0.003236246 | 3 | 0.02099 |
| **positive regulation of cell cycle arrest** | 2 | 0.003236246 | 3 | 0.02099 |
| **phosphatidylserine metabolic process** | 2 | 0.003236246 | 3 | 0.02099 |
| **phosphatidylserine biosynthetic process** | 2 | 0.003236246 | 3 | 0.02099 |
| **negative regulation of transport** | 2 | 0.003236246 | 3 | 0.02099 |

| | | | | |
|---|---|---|---|---|
| **negative regulation of ion transport** | 2 | 0.003236246 | 3 | 0.02099 |
| **menaquinone metabolic process** | 2 | 0.003236246 | 3 | 0.02099 |
| **menaquinone biosynthetic process** | 2 | 0.003236246 | 3 | 0.02099 |
| **bundle sheath cell fate specification** | 2 | 0.003236246 | 3 | 0.02099 |
| **activation of MAPK activity** | 2 | 0.003236246 | 3 | 0.02099 |
| **post-embryonic development** | 48 | 0.077669903 | 414 | 0.02127 |
| **nucleic acid-templated transcription** | 200 | 0.323624595 | 2036 | 0.02274 |
| **response to bacterium** | 14 | 0.022653722 | 91 | 0.02348 |
| **response to organic cyclic compound** | 16 | 0.025889968 | 109 | 0.02434 |
| **heterocycle biosynthetic process** | 240 | 0.388349515 | 2483 | 0.02518 |
| **regulation of cell cycle process** | 10 | 0.01618123 | 58 | 0.02528 |
| **phosphorelay signal transduction system** | 11 | 0.017799353 | 67 | 0.02726 |
| **trehalose biosynthetic process** | 6 | 0.009708738 | 28 | 0.0294 |
| **response to karrikin** | 5 | 0.008090615 | 21 | 0.02993 |
| **anatomical structure development** | 92 | 0.148867314 | 882 | 0.03055 |
| **system development** | 54 | 0.087378641 | 485 | 0.03067 |
| **transcription, DNA-templated** | 197 | 0.318770227 | 2022 | 0.03151 |
| **regulation of developmental growth** | 8 | 0.012944984 | 44 | 0.03241 |
| **regulation of innate immune response** | 6 | 0.009708738 | 29 | 0.03445 |
| **regulation of immune response** | 6 | 0.009708738 | 29 | 0.03445 |
| **stomatal complex morphogenesis** | 4 | 0.006472492 | 15 | 0.03471 |
| **phosphate ion transmembrane transport** | 3 | 0.004854369 | 9 | 0.03615 |
| **'de novo' posttranslational protein folding** | 3 | 0.004854369 | 9 | 0.03615 |
| **cellular response to alcohol** | 11 | 0.017799353 | 70 | 0.03633 |
| **cellular response to abscisic acid stimulus** | 11 | 0.017799353 | 70 | 0.03633 |
| **response to biotic stimulus** | 26 | 0.042071197 | 209 | 0.03673 |
| **cellular response to organic substance** | 43 | 0.069579288 | 378 | 0.03678 |
| **regulation of cellular metabolic process** | 230 | 0.372168285 | 2397 | 0.03701 |
| **response to auxin** | 23 | 0.037216828 | 181 | 0.03851 |
| **regulation of monopolar cell growth** | 2 | 0.003236246 | 4 | 0.03959 |
| **regulation of metal ion transport** | 2 | 0.003236246 | 4 | 0.03959 |
| **regulation of histone H3-K9 methylation** | 2 | 0.003236246 | 4 | 0.03959 |
| **monopolar cell growth** | 2 | 0.003236246 | 4 | 0.03959 |
| **mitotic G2 DNA damage checkpoint** | 2 | 0.003236246 | 4 | 0.03959 |
| **histone H3-K9 modification** | 2 | 0.003236246 | 4 | 0.03959 |
| **histone H3-K9 methylation** | 2 | 0.003236246 | 4 | 0.03959 |
| **G2 DNA damage checkpoint** | 2 | 0.003236246 | 4 | 0.03959 |
| **enzyme-directed rRNA pseudouridine** | 2 | 0.003236246 | 4 | 0.03959 |
| **chaperone-mediated protein transport** | 2 | 0.003236246 | 4 | 0.03959 |
| **cellular amide catabolic process** | 2 | 0.003236246 | 4 | 0.03959 |

| | | | | |
|---|---|---|---|---|
| **axis specification** | 2 | 0.003236246 | 4 | 0.03959 |
| **allantoin catabolic process** | 2 | 0.003236246 | 4 | 0.03959 |
| **regulation of primary metabolic process** | 229 | 0.370550162 | 2391 | 0.03995 |
| **trehalose metabolic process** | 6 | 0.009708738 | 30 | 0.04003 |
| **regulation of response to stress** | 16 | 0.025889968 | 116 | 0.04049 |
| **response to cold** | 14 | 0.022653722 | 98 | 0.04116 |
| **response to extracellular stimulus** | 10 | 0.01618123 | 63 | 0.04209 |
| **positive regulation of innate immune response** | 4 | 0.006472492 | 16 | 0.04321 |
| **positive regulation of immune system process** | 4 | 0.006472492 | 16 | 0.04321 |
| **positive regulation of immune response** | 4 | 0.006472492 | 16 | 0.04321 |
| **regulation of nitrogen compound metabolic** | 227 | 0.367313916 | 2375 | 0.0438 |
| **multicellular organismal process** | 98 | 0.158576052 | 963 | 0.04549 |
| **flower development** | 21 | 0.033980583 | 165 | 0.04572 |
| **glucan metabolic process** | 25 | 0.040453074 | 204 | 0.04642 |
| **defense response to bacterium** | 12 | 0.019417476 | 82 | 0.04786 |
| **regulation of microtubule-based process** | 3 | 0.004854369 | 10 | 0.0484 |
| **regulation of histone methylation** | 3 | 0.004854369 | 10 | 0.0484 |
| **mitotic DNA integrity checkpoint** | 3 | 0.004854369 | 10 | 0.0484 |
| **'de novo' pyrimidine nucleobase biosynthetic** | 3 | 0.004854369 | 10 | 0.0484 |
| **cellular polysaccharide metabolic process** | 28 | 0.045307443 | 235 | 0.04975 |

**Supplementary Table 6. Gene ontology enrichment of the 2707 genes differentially expressed between the *Pi-ta*$^-$ (X1) and *Pi-ta*$^{WT}$ (X7) lines.** *Pi-ta*$^-$ and *Pi-ta*$^{WT}$ lines are the *Pi-ta* knockout lines and the wild type line respectively. GO annotations are from experimentally verified datasets and differentially expressed genes were compared to the entire annotated *Oryza sativa* genome. Percentage is the number of genes enriched in one GO pathway divided the total number of genes differentially expressed between the *Pi-ta*$^-$ and *Pi-ta*$^{WT}$ lines.

| GO annotation | # DE in *Pi-ta*$^-$ vs *Pi-ta*$^{WT}$ | Percentage | # in Genome | *P* value |
|---|---|---|---|---|
| **cell cycle** | 47 | 0.017362394 | 395 | 2.60E-09 |
| **microtubule-based process** | 26 | 0.009604728 | 174 | 1.20E-07 |
| **cell cycle process** | 30 | 0.011082379 | 227 | 2.10E-07 |
| **cell division** | 24 | 0.008865903 | 165 | 6.00E-07 |
| **mitotic cell cycle process** | 20 | 0.007388253 | 122 | 7.60E-07 |
| **microtubule-based movement** | 17 | 0.006280015 | 93 | 1.10E-06 |
| **DNA conformation change** | 22 | 0.008127078 | 148 | 1.20E-06 |
| **movement of cell or subcellular** | 17 | 0.006280015 | 98 | 2.30E-06 |
| **chromosome organization** | 46 | 0.016992981 | 483 | 2.80E-06 |
| **organelle organization** | 90 | 0.033247137 | 1203 | 3.30E-06 |
| **mitotic cell cycle** | 23 | 0.008496491 | 173 | 5.00E-06 |
| **cytokinesis** | 10 | 0.003694126 | 45 | 3.10E-05 |
| **cellular component organization** | 130 | 0.048023642 | 2011 | 3.70E-05 |
| **DNA replication initiation** | 7 | 0.002585888 | 23 | 5.60E-05 |
| **mitotic cytokinesis** | 8 | 0.002955301 | 31 | 6.10E-05 |
| **cytoskeleton-dependent cytokinesis** | 8 | 0.002955301 | 32 | 7.90E-05 |
| **positive regulation of histone** | 3 | 0.001108238 | 3 | 9.80E-05 |
| **DNA geometric change** | 12 | 0.004432952 | 77 | 0.0002 |
| **DNA duplex unwinding** | 12 | 0.004432952 | 77 | 0.0002 |
| **response to water** | 14 | 0.005171777 | 101 | 0.00022 |
| **DNA replication** | 16 | 0.005910602 | 127 | 0.00025 |
| **DNA-dependent DNA replication** | 12 | 0.004432952 | 79 | 0.00026 |
| **positive regulation of chromosome** | 4 | 0.001477651 | 8 | 0.00027 |
| **cytokinetic process** | 5 | 0.001847063 | 15 | 0.00042 |
| **mitotic cytokinetic process** | 5 | 0.001847063 | 15 | 0.00042 |
| **cellular component organization or** | 139 | 0.051348356 | 2309 | 0.00051 |
| **response to water deprivation** | 13 | 0.004802364 | 98 | 0.00057 |
| **vesicle docking** | 8 | 0.002955301 | 43 | 0.00069 |
| **nuclear chromosome segregation** | 11 | 0.004063539 | 76 | 0.00071 |

| | | | | |
|---|---|---|---|---|
| **sister chromatid segregation** | 9 | 0.003324714 | 54 | 0.00075 |
| **membrane docking** | 8 | 0.002955301 | 44 | 0.00081 |
| **organelle localization by membrane** | 8 | 0.002955301 | 44 | 0.00081 |
| **positive regulation of protein** | 11 | 0.004063539 | 79 | 0.00098 |
| **negative regulation of transcription,** | 10 | 0.003694126 | 68 | 0.00107 |
| **response to acid chemical** | 27 | 0.009974141 | 312 | 0.00135 |
| **chromosome segregation** | 11 | 0.004063539 | 83 | 0.00148 |
| **positive regulation of histone** | 3 | 0.001108238 | 6 | 0.00177 |
| **positive regulation of chromatin** | 3 | 0.001108238 | 6 | 0.00177 |
| **regulation of response to salt stress** | 5 | 0.001847063 | 20 | 0.00181 |
| **maintenance of chromatin silencing** | 2 | 0.000738825 | 2 | 0.00213 |
| **transposition, RNA-mediated** | 2 | 0.000738825 | 2 | 0.00213 |
| **maintenance of protein location in** | 2 | 0.000738825 | 2 | 0.00213 |
| **positive regulation of histone H3-K9** | 2 | 0.000738825 | 2 | 0.00213 |
| **regulation of jasmonic acid biosynthetic** | 2 | 0.000738825 | 2 | 0.00213 |
| **regulation of histone H4 acetylation** | 2 | 0.000738825 | 2 | 0.00213 |
| **negative regulation of histone H4** | 2 | 0.000738825 | 2 | 0.00213 |
| **DNA metabolic process** | 45 | 0.016623569 | 625 | 0.00218 |
| **regulation of response to osmotic stress** | 5 | 0.001847063 | 21 | 0.00228 |
| **cytokinesis by cell plate formation** | 4 | 0.001477651 | 13 | 0.00232 |
| **response to salt stress** | 14 | 0.005171777 | 129 | 0.00257 |
| **negative regulation of nucleobase-** | 12 | 0.004432952 | 102 | 0.0026 |
| **negative regulation of RNA biosynthetic** | 11 | 0.004063539 | 89 | 0.00262 |
| **negative regulation of nucleic acid-** | 11 | 0.004063539 | 89 | 0.00262 |
| **chaperone mediated protein folding** | 3 | 0.001108238 | 7 | 0.00299 |
| **negative regulation of RNA metabolic** | 11 | 0.004063539 | 92 | 0.0034 |
| **chromatin silencing** | 6 | 0.002216476 | 33 | 0.00364 |
| **cell wall biogenesis** | 13 | 0.004802364 | 121 | 0.00392 |
| **asymmetric cell division** | 3 | 0.001108238 | 8 | 0.00463 |
| **sodium ion import** | 3 | 0.001108238 | 8 | 0.00463 |
| **inorganic cation import across plasma** | 3 | 0.001108238 | 8 | 0.00463 |
| **sodium ion import across plasma** | 3 | 0.001108238 | 8 | 0.00463 |
| **inorganic ion import across plasma** | 3 | 0.001108238 | 8 | 0.00463 |
| **DNA packaging** | 8 | 0.002955301 | 59 | 0.00553 |
| **response to abiotic stimulus** | 42 | 0.015515331 | 604 | 0.00555 |
| **cell wall organization** | 29 | 0.010712966 | 380 | 0.00577 |
| **menaquinone metabolic process** | 2 | 0.000738825 | 3 | 0.00621 |
| **menaquinone biosynthetic process** | 2 | 0.000738825 | 3 | 0.00621 |
| **regulation of jasmonic acid metabolic** | 2 | 0.000738825 | 3 | 0.00621 |
| **bundle sheath cell fate specification** | 2 | 0.000738825 | 3 | 0.00621 |

| | | | | |
|---|---|---|---|---|
| **external encapsulating structure** | 30 | 0.011082379 | 399 | 0.00621 |
| **microsporogenesis** | 3 | 0.001108238 | 9 | 0.0067 |
| **positive regulation of protein** | 3 | 0.001108238 | 9 | 0.0067 |
| **'de novo' posttranslational protein** | 3 | 0.001108238 | 9 | 0.0067 |
| **positive regulation of ubiquitin-protein** | 3 | 0.001108238 | 9 | 0.0067 |
| **import across plasma membrane** | 3 | 0.001108238 | 9 | 0.0067 |
| **positive regulation of protein** | 3 | 0.001108238 | 9 | 0.0067 |
| **response to inorganic substance** | 20 | 0.007388253 | 236 | 0.00674 |
| **negative regulation of gene expression,** | 6 | 0.002216476 | 38 | 0.00745 |
| **response to osmotic stress** | 14 | 0.005171777 | 147 | 0.00822 |
| **positive regulation of response to salt** | 4 | 0.001477651 | 18 | 0.00825 |
| **regulation of mitotic cell cycle** | 10 | 0.003694126 | 90 | 0.00854 |
| **DNA repair** | 25 | 0.009235316 | 325 | 0.00901 |
| **maintenance of DNA methylation** | 3 | 0.001108238 | 10 | 0.00925 |
| **regulation of histone methylation** | 3 | 0.001108238 | 10 | 0.00925 |
| **organelle fusion** | 9 | 0.003324714 | 78 | 0.00969 |
| **positive regulation of protein metabolic** | 13 | 0.004802364 | 136 | 0.01033 |
| **mitotic sister chromatid segregation** | 6 | 0.002216476 | 41 | 0.0108 |
| **vesicle fusion** | 8 | 0.002955301 | 66 | 0.01081 |
| **response to oxygen-containing** | 31 | 0.011451792 | 435 | 0.01142 |
| **organelle membrane fusion** | 8 | 0.002955301 | 67 | 0.01179 |
| **allantoin catabolic process** | 2 | 0.000738825 | 4 | 0.01203 |
| **mitotic chromosome condensation** | 2 | 0.000738825 | 4 | 0.01203 |
| **chloroplast-nucleus signaling pathway** | 2 | 0.000738825 | 4 | 0.01203 |
| **cellular amide catabolic process** | 2 | 0.000738825 | 4 | 0.01203 |
| **histone H3-K9 methylation** | 2 | 0.000738825 | 4 | 0.01203 |
| **regulation of histone H3-K9** | 2 | 0.000738825 | 4 | 0.01203 |
| **histone H3-K9 modification** | 2 | 0.000738825 | 4 | 0.01203 |
| **response to chitin** | 3 | 0.001108238 | 11 | 0.01229 |
| **exocytosis** | 10 | 0.003694126 | 96 | 0.01318 |
| **DNA alkylation** | 4 | 0.001477651 | 21 | 0.01446 |
| **DNA methylation** | 4 | 0.001477651 | 21 | 0.01446 |
| **chromosome separation** | 5 | 0.001847063 | 32 | 0.01487 |
| **response to cold** | 10 | 0.003694126 | 98 | 0.01508 |
| **regulation of developmental growth** | 6 | 0.002216476 | 44 | 0.0151 |
| **cold acclimation** | 3 | 0.001108238 | 12 | 0.01583 |
| **jasmonic acid biosynthetic process** | 3 | 0.001108238 | 12 | 0.01583 |
| **cellulose microfibril organization** | 3 | 0.001108238 | 12 | 0.01583 |
| **sexual sporulation** | 3 | 0.001108238 | 12 | 0.01583 |
| **sporulation** | 3 | 0.001108238 | 12 | 0.01583 |

| | | | | |
|---|---|---|---|---|
| **plant-type sporogenesis** | 3 | 0.001108238 | 12 | 0.01583 |
| **regulation of ubiquitin-protein** | 3 | 0.001108238 | 12 | 0.01583 |
| **cell wall assembly** | 3 | 0.001108238 | 12 | 0.01583 |
| **plant-type cell wall assembly** | 3 | 0.001108238 | 12 | 0.01583 |
| **meiotic cell cycle process** | 8 | 0.002955301 | 71 | 0.0164 |
| **meiotic cell cycle** | 9 | 0.003324714 | 85 | 0.01648 |
| **cellulose biosynthetic process** | 6 | 0.002216476 | 45 | 0.01676 |
| **regulation of cell cycle process** | 7 | 0.002585888 | 58 | 0.01701 |
| **cellular response to DNA damage** | 25 | 0.009235316 | 344 | 0.01736 |
| **organelle localization** | 10 | 0.003694126 | 101 | 0.01831 |
| **allantoin metabolic process** | 2 | 0.000738825 | 5 | 0.01944 |
| **methylation-dependent chromatin** | 2 | 0.000738825 | 5 | 0.01944 |
| **regulation of gene expression by genetic** | 2 | 0.000738825 | 5 | 0.01944 |
| **pyruvate transport** | 2 | 0.000738825 | 5 | 0.01944 |
| **mitochondrial pyruvate** | 2 | 0.000738825 | 5 | 0.01944 |
| **cellularization** | 2 | 0.000738825 | 5 | 0.01944 |
| **radial pattern formation** | 2 | 0.000738825 | 5 | 0.01944 |
| **negative regulation of histone** | 2 | 0.000738825 | 5 | 0.01944 |
| **response to decreased oxygen levels** | 2 | 0.000738825 | 5 | 0.01944 |
| **regulation of fatty acid biosynthetic** | 2 | 0.000738825 | 5 | 0.01944 |
| **regulation of DNA methylation** | 2 | 0.000738825 | 5 | 0.01944 |
| **regulation of seed growth** | 2 | 0.000738825 | 5 | 0.01944 |
| **pyruvate transmembrane transport** | 2 | 0.000738825 | 5 | 0.01944 |
| **negative regulation of protein** | 2 | 0.000738825 | 5 | 0.01944 |
| **negative regulation of peptidyl-lysine** | 2 | 0.000738825 | 5 | 0.01944 |
| **branched-chain amino acid biosynthetic** | 4 | 0.001477651 | 23 | 0.0199 |
| **nucleosome assembly** | 6 | 0.002216476 | 47 | 0.02045 |
| **cellular amino acid catabolic process** | 6 | 0.002216476 | 47 | 0.02045 |
| **positive regulation of transferase** | 8 | 0.002955301 | 74 | 0.02061 |
| **cellular response to stress** | 39 | 0.014407093 | 602 | 0.02122 |
| **positive regulation of cellular protein** | 12 | 0.004432952 | 135 | 0.02259 |
| **positive regulation of organelle** | 4 | 0.001477651 | 24 | 0.02302 |
| **membrane fusion** | 8 | 0.002955301 | 76 | 0.02381 |
| **chromatin organization** | 21 | 0.007757665 | 285 | 0.02414 |
| **regulation of histone modification** | 3 | 0.001108238 | 14 | 0.02445 |
| **regulation of protein ubiquitination** | 3 | 0.001108238 | 14 | 0.02445 |
| **purine-containing compound catabolic** | 3 | 0.001108238 | 14 | 0.02445 |
| **regulation of protein modification by** | 3 | 0.001108238 | 14 | 0.02445 |
| **mitochondrial transmembrane** | 3 | 0.001108238 | 14 | 0.02445 |
| **secretion by cell** | 10 | 0.003694126 | 106 | 0.02477 |

| | | | | |
|---|---|---|---|---|
| **response to chemical** | 68 | 0.025120059 | 1161 | 0.02535 |
| **response to organic substance** | 41 | 0.015145918 | 647 | 0.02543 |
| **beta-glucan biosynthetic process** | 7 | 0.002585888 | 63 | 0.02572 |
| **nuclear division** | 10 | 0.003694126 | 107 | 0.02624 |
| **response to salicylic acid** | 4 | 0.001477651 | 25 | 0.02643 |
| **xyloglucan metabolic process** | 6 | 0.002216476 | 50 | 0.02697 |
| **purine nucleobase catabolic process** | 2 | 0.000738825 | 6 | 0.02828 |
| **nucleoside triphosphate catabolic** | 2 | 0.000738825 | 6 | 0.02828 |
| **adaxial/abaxial pattern specification** | 2 | 0.000738825 | 6 | 0.02828 |
| **root meristem growth** | 2 | 0.000738825 | 6 | 0.02828 |
| **gas transport** | 2 | 0.000738825 | 6 | 0.02828 |
| **oxygen transport** | 2 | 0.000738825 | 6 | 0.02828 |
| **regulation of exocytosis** | 2 | 0.000738825 | 6 | 0.02828 |
| **regulation of fatty acid metabolic** | 2 | 0.000738825 | 6 | 0.02828 |
| **chromosome condensation** | 2 | 0.000738825 | 6 | 0.02828 |
| **mitochondrial respiratory chain** | 2 | 0.000738825 | 6 | 0.02828 |
| **regulation of histone acetylation** | 2 | 0.000738825 | 6 | 0.02828 |
| **histone H4 acetylation** | 2 | 0.000738825 | 6 | 0.02828 |
| **response to freezing** | 2 | 0.000738825 | 6 | 0.02828 |
| **regulation of secretion** | 2 | 0.000738825 | 6 | 0.02828 |
| **regulation of mitotic spindle** | 2 | 0.000738825 | 6 | 0.02828 |
| **response to oxygen levels** | 2 | 0.000738825 | 6 | 0.02828 |
| **genetic imprinting** | 2 | 0.000738825 | 6 | 0.02828 |
| **seed growth** | 2 | 0.000738825 | 6 | 0.02828 |
| **regulation of chlorophyll metabolic** | 2 | 0.000738825 | 6 | 0.02828 |
| **regulation of spindle organization** | 2 | 0.000738825 | 6 | 0.02828 |
| **regulation of protein acetylation** | 2 | 0.000738825 | 6 | 0.02828 |
| **regulation of secretion by cell** | 2 | 0.000738825 | 6 | 0.02828 |
| **positive regulation of ubiquitin protein** | 2 | 0.000738825 | 6 | 0.02828 |
| **regulation of peptidyl-lysine acetylation** | 2 | 0.000738825 | 6 | 0.02828 |
| **organelle fission** | 12 | 0.004432952 | 140 | 0.02899 |
| **postreplication repair** | 3 | 0.001108238 | 15 | 0.02954 |
| **regulation of lipid biosynthetic process** | 3 | 0.001108238 | 15 | 0.02954 |
| **mitotic cell cycle phase transition** | 5 | 0.001847063 | 38 | 0.02959 |
| **response to temperature stimulus** | 14 | 0.005171777 | 173 | 0.02995 |
| **megagametogenesis** | 4 | 0.001477651 | 26 | 0.03013 |
| **cell cycle phase transition** | 5 | 0.001847063 | 39 | 0.03269 |
| **cytoskeleton organization** | 13 | 0.004802364 | 159 | 0.03312 |
| **cell wall organization or biogenesis** | 34 | 0.01256003 | 530 | 0.03381 |
| **DNA methylation or demethylation** | 4 | 0.001477651 | 27 | 0.03412 |

| | | | | |
|---|---|---|---|---|
| **secretion** | 10 | 0.003694126 | 112 | 0.03452 |
| **microtubule cytoskeleton organization** | 7 | 0.002585888 | 67 | 0.03458 |
| **regulation of protein modification** | 13 | 0.004802364 | 160 | 0.03459 |
| **vacuolar transport** | 6 | 0.002216476 | 53 | 0.03473 |
| **chromatin assembly** | 6 | 0.002216476 | 53 | 0.03473 |
| **nucleosome organization** | 6 | 0.002216476 | 53 | 0.03473 |
| **mitotic nuclear division** | 6 | 0.002216476 | 53 | 0.03473 |
| **response to lipid** | 17 | 0.006280015 | 227 | 0.03473 |
| **plant-type primary cell wall biogenesis** | 3 | 0.001108238 | 16 | 0.03513 |
| **negative regulation of nitrogen** | 20 | 0.007388253 | 280 | 0.03614 |
| **G1/S transition of mitotic cell cycle** | 2 | 0.000738825 | 7 | 0.03839 |
| **glycine catabolic process** | 2 | 0.000738825 | 7 | 0.03839 |
| **deoxyribonucleotide biosynthetic** | 2 | 0.000738825 | 7 | 0.03839 |
| **genetic transfer** | 2 | 0.000738825 | 7 | 0.03839 |
| **DNA mediated transformation** | 2 | 0.000738825 | 7 | 0.03839 |
| **poly(A)+ mRNA export from nucleus** | 2 | 0.000738825 | 7 | 0.03839 |
| **negative regulation of histone** | 2 | 0.000738825 | 7 | 0.03839 |
| **UDP-L-arabinose metabolic process** | 2 | 0.000738825 | 7 | 0.03839 |
| **multi-organism cellular process** | 2 | 0.000738825 | 7 | 0.03839 |
| **cell cycle G1/S phase transition** | 2 | 0.000738825 | 7 | 0.03839 |
| **regulation of microtubule cytoskeleton** | 2 | 0.000738825 | 7 | 0.03839 |
| **mitochondrial respiratory chain** | 2 | 0.000738825 | 7 | 0.03839 |
| **nucleoside phosphate catabolic process** | 2 | 0.000738825 | 7 | 0.03839 |
| **branched-chain amino acid metabolic** | 4 | 0.001477651 | 28 | 0.0384 |
| **response to topologically incorrect** | 4 | 0.001477651 | 28 | 0.0384 |
| **response to stimulus** | 175 | 0.064647211 | 3356 | 0.03876 |
| **vesicle organization** | 8 | 0.002955301 | 84 | 0.0401 |
| **jasmonic acid metabolic process** | 3 | 0.001108238 | 17 | 0.04123 |
| **cellular glucan metabolic process** | 15 | 0.00554119 | 198 | 0.04168 |
| **DNA modification** | 4 | 0.001477651 | 29 | 0.04297 |
| **cellular polysaccharide metabolic** | 17 | 0.006280015 | 235 | 0.04573 |
| **activation of MAPK activity involved in** | 1 | 0.000369413 | 1 | 0.04623 |
| **photoreactive repair** | 1 | 0.000369413 | 1 | 0.04623 |
| **assembly of actomyosin apparatus** | 1 | 0.000369413 | 1 | 0.04623 |
| **phragmoplast assembly** | 1 | 0.000369413 | 1 | 0.04623 |
| **action potential** | 1 | 0.000369413 | 1 | 0.04623 |
| **dUMP biosynthetic process** | 1 | 0.000369413 | 1 | 0.04623 |
| **pyrimidine nucleotide catabolic process** | 1 | 0.000369413 | 1 | 0.04623 |
| **DNA unwinding involved in DNA** | 1 | 0.000369413 | 1 | 0.04623 |
| **osmosensory signaling pathway** | 1 | 0.000369413 | 1 | 0.04623 |

| | | | | |
|---|---|---|---|---|
| **pyrimidine nucleoside triphosphate** | 1 | 0.000369413 | 1 | 0.04623 |
| **pyrimidine deoxyribonucleoside** | 1 | 0.000369413 | 1 | 0.04623 |
| **pyrimidine deoxyribonucleotide** | 1 | 0.000369413 | 1 | 0.04623 |
| **radial microtubular system formation** | 1 | 0.000369413 | 1 | 0.04623 |
| **endosperm cellularization** | 1 | 0.000369413 | 1 | 0.04623 |
| **vegetative meristem growth** | 1 | 0.000369413 | 1 | 0.04623 |
| **magnesium ion homeostasis** | 1 | 0.000369413 | 1 | 0.04623 |
| **mitochondrial DNA metabolic process** | 1 | 0.000369413 | 1 | 0.04623 |
| **chlorophyll cycle** | 1 | 0.000369413 | 1 | 0.04623 |
| **DNA rewinding** | 1 | 0.000369413 | 1 | 0.04623 |
| **dUMP metabolic process** | 1 | 0.000369413 | 1 | 0.04623 |
| **dUTP metabolic process** | 1 | 0.000369413 | 1 | 0.04623 |
| **dUTP catabolic process** | 1 | 0.000369413 | 1 | 0.04623 |
| **membrane depolarization** | 1 | 0.000369413 | 1 | 0.04623 |
| **positive regulation of histone H3-K27** | 1 | 0.000369413 | 1 | 0.04623 |
| **pri-miRNA transcription from RNA** | 1 | 0.000369413 | 1 | 0.04623 |
| **membrane depolarization during action** | 1 | 0.000369413 | 1 | 0.04623 |
| **tubulin deacetylation** | 1 | 0.000369413 | 1 | 0.04623 |
| **nicotinate metabolic process** | 1 | 0.000369413 | 1 | 0.04623 |
| **assembly of actomyosin apparatus** | 1 | 0.000369413 | 1 | 0.04623 |
| **positive regulation of defense response** | 1 | 0.000369413 | 1 | 0.04623 |
| **polyuridylation-dependent mRNA** | 1 | 0.000369413 | 1 | 0.04623 |
| **malonyl-CoA biosynthetic process** | 1 | 0.000369413 | 1 | 0.04623 |
| **galactose metabolic process** | 3 | 0.001108238 | 18 | 0.04783 |
| **purine nucleobase metabolic process** | 3 | 0.001108238 | 18 | 0.04783 |
| **maintenance of location in cell** | 3 | 0.001108238 | 18 | 0.04783 |
| **G-protein coupled receptor signaling** | 4 | 0.001477651 | 30 | 0.04783 |
| **cellulose metabolic process** | 7 | 0.002585888 | 72 | 0.04823 |
| **gametophyte development** | 10 | 0.003694126 | 119 | 0.04894 |
| **membrane organization** | 12 | 0.004432952 | 152 | 0.04948 |
| **respiratory chain complex IV assembly** | 2 | 0.000738825 | 8 | 0.04964 |
| **serine family amino acid catabolic** | 2 | 0.000738825 | 8 | 0.04964 |
| **nucleoside catabolic process** | 2 | 0.000738825 | 8 | 0.04964 |
| **DNA methylation on cytosine** | 2 | 0.000738825 | 8 | 0.04964 |
| **pyrimidine-containing compound** | 2 | 0.000738825 | 8 | 0.04964 |
| **maintenance of protein localization in** | 2 | 0.000738825 | 8 | 0.04964 |
| **C-5 methylation of cytosine** | 2 | 0.000738825 | 8 | 0.04964 |
| **regulation of tetrapyrrole metabolic** | 2 | 0.000738825 | 8 | 0.04964 |
| **glycosyl compound catabolic process** | 2 | 0.000738825 | 8 | 0.04964 |

**Supplementary Table 7. Gene ontology enrichment of the 2901 genes differentially expressed between the *Pi-ta*⁻ (X4) and *Pi-ta*$^{WT}$ (X7) lines.** *Pi-ta*$^{-}$ and *Pi-ta*$^{WT}$ lines are the *Pi-ta* knockout lines and the wild type line respectively. GO annotations are from experimentally verified datasets and differentially expressed genes were compared to the entire annotated *Oryza sativa* genome. Percentage is the number of genes enriched in one GO pathway divided the total number of genes differentially expressed between the *Pi-ta*$^{-}$ and *Pi-ta*$^{WT}$ lines.

| GO annotation | # DE in *Pi-ta*$^{-}$ vs *Pi-ta*$^{WT}$ | Percentage | # in Genome | *P* value |
|---|---|---|---|---|
| **response to stimulus** | 383 | 0.13202344 | 3356 | 2.10E-06 |
| **post-embryonic plant morphogenesis** | 19 | 0.006549466 | 66 | 5.40E-06 |
| **response to abiotic stimulus** | 89 | 0.030679076 | 604 | 7.60E-06 |
| **asymmetric cell division** | 6 | 0.002068252 | 8 | 1.50E-05 |
| **stomatal complex morphogenesis** | 8 | 0.00275767 | 15 | 1.90E-05 |
| **anatomical structure development** | 118 | 0.040675629 | 882 | 2.80E-05 |
| **multicellular organism development** | 107 | 0.036883833 | 792 | 4.50E-05 |
| **guard cell differentiation** | 7 | 0.002412961 | 13 | 6.10E-05 |
| **stomatal complex development** | 8 | 0.00275767 | 18 | 0.0001 |
| **chaperone mediated protein folding** | 5 | 0.001723544 | 7 | 0.00012 |
| **DNA conformation change** | 28 | 0.009651844 | 148 | 0.00021 |
| **response to water** | 21 | 0.007238883 | 101 | 0.00034 |
| **xylem development** | 4 | 0.001378835 | 5 | 0.00034 |
| **response to external stimulus** | 46 | 0.015856601 | 296 | 0.00036 |
| **cold acclimation** | 6 | 0.002068252 | 12 | 0.00036 |
| **pattern specification process** | 13 | 0.004481213 | 50 | 0.00049 |
| **regionalization** | 12 | 0.004136505 | 44 | 0.0005 |
| **response to extracellular stimulus** | 15 | 0.005170631 | 63 | 0.00052 |
| **response to nutrient levels** | 14 | 0.004825922 | 57 | 0.00057 |
| **response to water deprivation** | 20 | 0.006894174 | 98 | 0.0006 |
| **post-embryonic development** | 59 | 0.020337815 | 414 | 0.0006 |
| **anion homeostasis** | 6 | 0.002068252 | 13 | 0.00062 |
| **'de novo' posttranslational protein** | 5 | 0.001723544 | 9 | 0.00063 |
| **multicellular organismal process** | 119 | 0.041020338 | 963 | 0.00069 |
| **microtubule-based movement** | 19 | 0.006549466 | 93 | 0.0008 |
| **developmental process** | 132 | 0.045501551 | 1090 | 0.00081 |
| **response to chemical** | 139 | 0.047914512 | 1161 | 0.00095 |
| **phloem or xylem histogenesis** | 6 | 0.002068252 | 14 | 0.00099 |
| **flavonoid biosynthetic process** | 8 | 0.00275767 | 24 | 0.00103 |

| | | | | |
|---|---|---|---|---|
| **response to acid chemical** | 46 | 0.015856601 | 312 | 0.00114 |
| **plant epidermis morphogenesis** | 9 | 0.003102378 | 30 | 0.00119 |
| **shoot system development** | 40 | 0.013788349 | 263 | 0.00128 |
| **response to starvation** | 12 | 0.004136505 | 49 | 0.00142 |
| **movement of cell or subcellular** | 19 | 0.006549466 | 98 | 0.00154 |
| **DNA geometric change** | 16 | 0.00551534 | 77 | 0.00166 |
| **DNA duplex unwinding** | 16 | 0.00551534 | 77 | 0.00166 |
| **carbohydrate metabolic process** | 120 | 0.041365047 | 998 | 0.00179 |
| **response to oxygen-containing** | 59 | 0.020337815 | 435 | 0.002 |
| **shoot system morphogenesis** | 12 | 0.004136505 | 51 | 0.00205 |
| **response to organic substance** | 82 | 0.028266115 | 647 | 0.00228 |
| **tissue development** | 23 | 0.007928301 | 132 | 0.00233 |
| **cell cycle** | 54 | 0.018614271 | 395 | 0.00255 |
| **intracellular signal transduction** | 54 | 0.018614271 | 395 | 0.00255 |
| **regulation of metal ion transport** | 3 | 0.001034126 | 4 | 0.00298 |
| **photomorphogenesis** | 8 | 0.00275767 | 28 | 0.00312 |
| **flavonoid metabolic process** | 8 | 0.00275767 | 28 | 0.00312 |
| **peptidyl-histidine phosphorylation** | 5 | 0.001723544 | 12 | 0.00312 |
| **cellular response to hormone** | 45 | 0.015511892 | 320 | 0.00327 |
| **response to abscisic acid** | 25 | 0.008617718 | 152 | 0.00347 |
| **regulation of nucleobase-containing** | 217 | 0.074801792 | 1973 | 0.00361 |
| **DNA replication initiation** | 7 | 0.002412961 | 23 | 0.00381 |
| **positive regulation of chromosome** | 4 | 0.001378835 | 8 | 0.00383 |
| **regulation of RNA metabolic process** | 213 | 0.073422958 | 1937 | 0.00397 |
| **system development** | 63 | 0.021716649 | 485 | 0.00401 |
| **response to hormone** | 67 | 0.023095484 | 522 | 0.00408 |
| **cellular response to endogenous** | 45 | 0.015511892 | 324 | 0.00414 |
| **auxin-activated signaling pathway** | 18 | 0.006204757 | 99 | 0.00418 |
| **cellular response to phosphate** | 6 | 0.002068252 | 18 | 0.00444 |
| **cellular response to extracellular** | 11 | 0.003791796 | 49 | 0.00458 |
| **cell wall organization** | 51 | 0.017580145 | 380 | 0.00479 |
| **anatomical structure morphogenesis** | 34 | 0.011720097 | 231 | 0.00482 |
| **response to endogenous stimulus** | 67 | 0.023095484 | 526 | 0.00487 |
| **response to alcohol** | 25 | 0.008617718 | 156 | 0.0049 |
| **plant epidermal cell differentiation** | 8 | 0.00275767 | 30 | 0.00497 |
| **cellular response to auxin stimulus** | 18 | 0.006204757 | 101 | 0.00519 |
| **cell division** | 26 | 0.008962427 | 165 | 0.00524 |
| **regulation of transcription, DNA-** | 207 | 0.071354705 | 1889 | 0.00527 |
| **cellular response to external** | 11 | 0.003791796 | 50 | 0.00539 |
| **lipid transport** | 17 | 0.005860048 | 94 | 0.00558 |

| | | | | |
|---|---|---|---|---|
| **regulation of nucleic acid-templated** | 208 | 0.071699414 | 1902 | 0.00564 |
| **regulation of RNA biosynthetic** | 208 | 0.071699414 | 1902 | 0.00564 |
| **hormone-mediated signaling** | 43 | 0.014822475 | 313 | 0.00604 |
| **cellular response to nutrient levels** | 10 | 0.003447087 | 44 | 0.00614 |
| **regulation of meristem growth** | 4 | 0.001378835 | 9 | 0.00639 |
| **phosphate ion transmembrane** | 4 | 0.001378835 | 9 | 0.00639 |
| **cellular carbohydrate metabolic** | 47 | 0.01620131 | 350 | 0.00651 |
| **response to lipid** | 33 | 0.011375388 | 227 | 0.00655 |
| **response to inorganic substance** | 34 | 0.011720097 | 236 | 0.00672 |
| **glycosylceramide metabolic process** | 3 | 0.001034126 | 5 | 0.00694 |
| **glucosylceramide metabolic process** | 3 | 0.001034126 | 5 | 0.00694 |
| **glycosphingolipid metabolic process** | 3 | 0.001034126 | 5 | 0.00694 |
| **cell proliferation** | 8 | 0.00275767 | 32 | 0.00756 |
| **cellular response to starvation** | 9 | 0.003102378 | 39 | 0.00824 |
| **cell wall organization or biogenesis** | 66 | 0.022750776 | 530 | 0.00855 |
| **regulation of chlorophyll catabolic** | 2 | 0.000689417 | 2 | 0.00863 |
| **regulation of iron ion transport** | 2 | 0.000689417 | 2 | 0.00863 |
| **negative regulation of iron ion** | 2 | 0.000689417 | 2 | 0.00863 |
| **pigmentation** | 2 | 0.000689417 | 2 | 0.00863 |
| **pigment accumulation** | 2 | 0.000689417 | 2 | 0.00863 |
| **pigment accumulation in response to** | 2 | 0.000689417 | 2 | 0.00863 |
| **pigment accumulation in tissues in** | 2 | 0.000689417 | 2 | 0.00863 |
| **pigment accumulation in tissues** | 2 | 0.000689417 | 2 | 0.00863 |
| **anthocyanin accumulation in tissues** | 2 | 0.000689417 | 2 | 0.00863 |
| **maintenance of protein location in** | 2 | 0.000689417 | 2 | 0.00863 |
| **secondary growth** | 2 | 0.000689417 | 2 | 0.00863 |
| **lateral growth** | 2 | 0.000689417 | 2 | 0.00863 |
| **regulation of tetrapyrrole catabolic** | 2 | 0.000689417 | 2 | 0.00863 |
| **lipid localization** | 18 | 0.006204757 | 106 | 0.00864 |
| **peptidyl-histidine modification** | 5 | 0.001723544 | 15 | 0.00935 |
| **cell wall modification** | 11 | 0.003791796 | 54 | 0.0098 |
| **transcription, DNA-templated** | 217 | 0.074801792 | 2022 | 0.01068 |
| **nucleic acid-templated transcription** | 218 | 0.075146501 | 2036 | 0.01156 |
| **external encapsulating structure** | 51 | 0.017580145 | 399 | 0.01205 |
| **RNA biosynthetic process** | 218 | 0.075146501 | 2039 | 0.01229 |
| **GDP-mannose biosynthetic process** | 3 | 0.001034126 | 6 | 0.01292 |
| **positive regulation of histone** | 3 | 0.001034126 | 6 | 0.01292 |
| **phosphate ion homeostasis** | 3 | 0.001034126 | 6 | 0.01292 |
| **divalent inorganic anion homeostasis** | 3 | 0.001034126 | 6 | 0.01292 |
| **trivalent inorganic anion** | 3 | 0.001034126 | 6 | 0.01292 |

| | | | | |
|---|---|---|---|---|
| **positive regulation of chromatin** | 3 | 0.001034126 | 6 | 0.01292 |
| **RNA destabilization** | 4 | 0.001378835 | 11 | 0.01438 |
| **mRNA destabilization** | 4 | 0.001378835 | 11 | 0.01438 |
| **3'-UTR-mediated mRNA** | 4 | 0.001378835 | 11 | 0.01438 |
| **'de novo' protein folding** | 7 | 0.002412961 | 29 | 0.01485 |
| **regulation of cellular metabolic** | 252 | 0.086866598 | 2397 | 0.01542 |
| **signal transduction** | 160 | 0.055153395 | 1466 | 0.01562 |
| **cell communication** | 176 | 0.060668735 | 1627 | 0.01578 |
| **biological regulation** | 483 | 0.166494312 | 4798 | 0.01592 |
| **cell wall biogenesis** | 19 | 0.006549466 | 121 | 0.016 |
| **cellular response to organic** | 48 | 0.016546019 | 378 | 0.01622 |
| **signaling** | 160 | 0.055153395 | 1468 | 0.01634 |
| **nucleobase-containing compound** | 244 | 0.084108928 | 2319 | 0.01651 |
| **response to osmotic stress** | 22 | 0.007583592 | 147 | 0.01713 |
| **chaperone-mediated protein folding** | 10 | 0.003447087 | 51 | 0.01748 |
| **response to cold** | 16 | 0.00551534 | 98 | 0.01827 |
| **inorganic anion transmembrane** | 8 | 0.00275767 | 37 | 0.01835 |
| **polysaccharide metabolic process** | 46 | 0.015856601 | 364 | 0.01976 |
| **regulation of ion transport** | 6 | 0.002068252 | 24 | 0.01989 |
| **plant epidermis development** | 10 | 0.003447087 | 52 | 0.0199 |
| **cellular response to stimulus** | 237 | 0.081695967 | 2258 | 0.01998 |
| **lateral root formation** | 4 | 0.001378835 | 12 | 0.02002 |
| **meristem growth** | 4 | 0.001378835 | 12 | 0.02002 |
| **post-embryonic plant organ** | 9 | 0.003102378 | 45 | 0.02086 |
| **G1/S transition of mitotic cell cycle** | 3 | 0.001034126 | 7 | 0.02106 |
| **GDP-mannose metabolic process** | 3 | 0.001034126 | 7 | 0.02106 |
| **cell cycle G1/S phase transition** | 3 | 0.001034126 | 7 | 0.02106 |
| **regulation of meristem development** | 7 | 0.002412961 | 31 | 0.02126 |
| **fruit development** | 21 | 0.007238883 | 143 | 0.02366 |
| **glycyl-tRNA aminoacylation** | 2 | 0.000689417 | 3 | 0.02429 |
| **menaquinone metabolic process** | 2 | 0.000689417 | 3 | 0.02429 |
| **menaquinone biosynthetic process** | 2 | 0.000689417 | 3 | 0.02429 |
| **positive regulation of histone** | 2 | 0.000689417 | 3 | 0.02429 |
| **amino acid export** | 2 | 0.000689417 | 3 | 0.02429 |
| **negative regulation of ion transport** | 2 | 0.000689417 | 3 | 0.02429 |
| **protein import into mitochondrial** | 2 | 0.000689417 | 3 | 0.02429 |
| **negative regulation of transport** | 2 | 0.000689417 | 3 | 0.02429 |
| **amino acid homeostasis** | 2 | 0.000689417 | 3 | 0.02429 |
| **bundle sheath cell fate specification** | 2 | 0.000689417 | 3 | 0.02429 |
| **regulation of cellular response to** | 2 | 0.000689417 | 3 | 0.02429 |

| | | | | |
|---|---|---|---|---|
| **positive regulation of cellular** | 2 | 0.000689417 | 3 | 0.02429 |
| **positive regulation of response to** | 2 | 0.000689417 | 3 | 0.02429 |
| **regulation of superoxide dismutase** | 2 | 0.000689417 | 3 | 0.02429 |
| **positive regulation of superoxide** | 2 | 0.000689417 | 3 | 0.02429 |
| **positive regulation of removal of** | 2 | 0.000689417 | 3 | 0.02429 |
| **regulation of removal of superoxide** | 2 | 0.000689417 | 3 | 0.02429 |
| **cellular lipid catabolic process** | 10 | 0.003447087 | 54 | 0.02545 |
| **regulation of cellular macromolecule** | 215 | 0.074112375 | 2047 | 0.02559 |
| **negative regulation of catalytic** | 25 | 0.008617718 | 179 | 0.02571 |
| **regulation of nitrogen compound** | 247 | 0.085143054 | 2375 | 0.02571 |
| **response to stress** | 188 | 0.06480524 | 1773 | 0.02607 |
| **seed development** | 20 | 0.006894174 | 136 | 0.0264 |
| **response to unfolded protein** | 5 | 0.001723544 | 19 | 0.02652 |
| **regulation of cell proliferation** | 5 | 0.001723544 | 19 | 0.02652 |
| **protein phosphorylation** | 173 | 0.059634609 | 1622 | 0.02662 |
| **cellular iron ion homeostasis** | 4 | 0.001378835 | 13 | 0.02683 |
| **gravitropism** | 4 | 0.001378835 | 13 | 0.02683 |
| **lateral root morphogenesis** | 4 | 0.001378835 | 13 | 0.02683 |
| **protein refolding** | 4 | 0.001378835 | 13 | 0.02683 |
| **regulation of mRNA stability** | 4 | 0.001378835 | 13 | 0.02683 |
| **positive regulation of mRNA** | 4 | 0.001378835 | 13 | 0.02683 |
| **positive regulation of mRNA** | 4 | 0.001378835 | 13 | 0.02683 |
| **response to red or far red light** | 12 | 0.004136505 | 70 | 0.02684 |
| **cellular response to abscisic acid** | 12 | 0.004136505 | 70 | 0.02684 |
| **cellular response to alcohol** | 12 | 0.004136505 | 70 | 0.02684 |
| **plant organ formation** | 9 | 0.003102378 | 47 | 0.02716 |
| **regulation of primary metabolic** | 248 | 0.085487763 | 2391 | 0.02828 |
| **regulation of macromolecule** | 216 | 0.074457084 | 2064 | 0.02887 |
| **negative regulation of molecular** | 25 | 0.008617718 | 181 | 0.029 |
| **peptidyl-threonine phosphorylation** | 6 | 0.002068252 | 26 | 0.02902 |
| **peptidyl-threonine modification** | 6 | 0.002068252 | 26 | 0.02902 |
| **regulation of cellular biosynthetic** | 218 | 0.075146501 | 2085 | 0.02914 |
| **response to salt stress** | 19 | 0.006549466 | 129 | 0.02948 |
| **cell wall macromolecule metabolic** | 19 | 0.006549466 | 129 | 0.02948 |
| **DNA-dependent DNA replication** | 13 | 0.004481213 | 79 | 0.02959 |
| **reproductive structure development** | 41 | 0.014133058 | 327 | 0.02971 |
| **reproductive system development** | 41 | 0.014133058 | 327 | 0.02971 |
| **phosphate-containing compound** | 280 | 0.096518442 | 2724 | 0.02971 |
| **regulation of biosynthetic process** | 218 | 0.075146501 | 2088 | 0.0307 |
| **phosphorus metabolic process** | 284 | 0.097897277 | 2768 | 0.03094 |

| | | | | |
|---|---|---|---|---|
| **response to biotic stimulus** | 28 | 0.009651844 | 209 | 0.03114 |
| **cinnamic acid biosynthetic process** | 3 | 0.001034126 | 8 | 0.03141 |
| **cinnamic acid metabolic process** | 3 | 0.001034126 | 8 | 0.03141 |
| **regulation of homeostatic process** | 3 | 0.001034126 | 8 | 0.03141 |
| **monovalent inorganic anion** | 3 | 0.001034126 | 8 | 0.03141 |
| **regulation of cellular process** | 396 | 0.136504654 | 3938 | 0.03255 |
| **negative regulation of peptidase** | 11 | 0.003791796 | 64 | 0.03261 |
| **negative regulation of endopeptidase** | 11 | 0.003791796 | 64 | 0.03261 |
| **negative regulation of proteolysis** | 11 | 0.003791796 | 64 | 0.03261 |
| **iron ion transport** | 5 | 0.001723544 | 20 | 0.03273 |
| **lateral root development** | 5 | 0.001723544 | 20 | 0.03273 |
| **developmental process involved in** | 45 | 0.015511892 | 367 | 0.0331 |
| **G2/M transition of mitotic cell cycle** | 4 | 0.001378835 | 14 | 0.03486 |
| **post-embryonic root morphogenesis** | 4 | 0.001378835 | 14 | 0.03486 |
| **regulation of G2/M transition of** | 4 | 0.001378835 | 14 | 0.03486 |
| **regulation of RNA stability** | 4 | 0.001378835 | 14 | 0.03486 |
| **phosphorylation** | 213 | 0.073422958 | 2045 | 0.03535 |
| **galacturonan metabolic process** | 12 | 0.004136505 | 73 | 0.03595 |
| **pectin metabolic process** | 12 | 0.004136505 | 73 | 0.03595 |
| **regulation of endopeptidase activity** | 11 | 0.003791796 | 65 | 0.03608 |
| **response to radiation** | 36 | 0.012409514 | 286 | 0.03765 |
| **xyloglucan metabolic process** | 9 | 0.003102378 | 50 | 0.03895 |
| **post-embryonic plant organ** | 7 | 0.002412961 | 35 | 0.03935 |
| **anatomical structure formation** | 12 | 0.004136505 | 74 | 0.03942 |
| **phospholipid dephosphorylation** | 5 | 0.001723544 | 21 | 0.03977 |
| **phosphatidylinositol** | 5 | 0.001723544 | 21 | 0.03977 |
| **response to karrikin** | 5 | 0.001723544 | 21 | 0.03977 |
| **regulation of peptidase activity** | 11 | 0.003791796 | 66 | 0.0398 |
| **trehalose biosynthetic process** | 6 | 0.002068252 | 28 | 0.04055 |
| **meristem maintenance** | 6 | 0.002068252 | 28 | 0.04055 |
| **response to topologically incorrect** | 6 | 0.002068252 | 28 | 0.04055 |
| **phosphorelay signal transduction** | 11 | 0.003791796 | 67 | 0.04378 |
| **isoprenoid catabolic process** | 4 | 0.001378835 | 15 | 0.04413 |
| **response to gravity** | 4 | 0.001378835 | 15 | 0.04413 |
| **cell cycle G2/M phase transition** | 4 | 0.001378835 | 15 | 0.04413 |
| **regulation of mRNA catabolic** | 4 | 0.001378835 | 15 | 0.04413 |
| **regulation of cell cycle G2/M phase** | 4 | 0.001378835 | 15 | 0.04413 |
| **aromatic compound biosynthetic** | 259 | 0.089279559 | 2533 | 0.04421 |
| **heterocycle biosynthetic process** | 254 | 0.087556015 | 2483 | 0.04537 |
| **glucosylceramide catabolic process** | 2 | 0.000689417 | 4 | 0.04561 |

| | | | | |
|---|---|---|---|---|
| **mitotic chromosome condensation** | 2 | 0.000689417 | 4 | 0.04561 |
| **mitotic G2 DNA damage checkpoint** | 2 | 0.000689417 | 4 | 0.04561 |
| **axis specification** | 2 | 0.000689417 | 4 | 0.04561 |
| **glycolipid catabolic process** | 2 | 0.000689417 | 4 | 0.04561 |
| **G2 DNA damage checkpoint** | 2 | 0.000689417 | 4 | 0.04561 |
| **monopolar cell growth** | 2 | 0.000689417 | 4 | 0.04561 |
| **amino acid import** | 2 | 0.000689417 | 4 | 0.04561 |
| **glycosylceramide catabolic process** | 2 | 0.000689417 | 4 | 0.04561 |
| **glycosphingolipid catabolic process** | 2 | 0.000689417 | 4 | 0.04561 |
| **ceramide catabolic process** | 2 | 0.000689417 | 4 | 0.04561 |
| **kinetochore assembly** | 2 | 0.000689417 | 4 | 0.04561 |
| **kinetochore organization** | 2 | 0.000689417 | 4 | 0.04561 |
| **regulation of monopolar cell growth** | 2 | 0.000689417 | 4 | 0.04561 |
| **regulation of response to reactive** | 2 | 0.000689417 | 4 | 0.04561 |
| **positive regulation of response to** | 2 | 0.000689417 | 4 | 0.04561 |
| **regulation of seed maturation** | 2 | 0.000689417 | 4 | 0.04561 |
| **DNA replication** | 18 | 0.006204757 | 127 | 0.04637 |
| **carbohydrate phosphorylation** | 6 | 0.002068252 | 29 | 0.04725 |
| **post-embryonic root development** | 5 | 0.001723544 | 22 | 0.04766 |
| **response to external biotic stimulus** | 25 | 0.008617718 | 190 | 0.04788 |
| **chromosome organization** | 56 | 0.019303688 | 483 | 0.04901 |
| **regulation of biological process** | 423 | 0.145811789 | 4255 | 0.04948 |

**Supplementary Table 8. Gene ontology enrichment of the 1443 genes differentially expressed between the *Pi-ta*$^-$ (X6) and *Pi-ta*$^{WT}$ (X7) lines.** *Pi-ta*$^-$ and *Pi-ta*$^{WT}$ lines are the *Pi-ta* knockout lines and the wild type line respectively. GO annotations are from experimentally verified datasets and differentially expressed genes were compared to the entire annotated *Oryza sativa* genome. Percentage is the number of genes enriched in one GO pathway divided the total number of genes differentially expressed between the *Pi-ta*$^-$ and *Pi-ta*$^{WT}$ lines.

| GO annotation | # DE in *Pi-ta*$^-$ vs *Pi-ta*$^{WT}$ | Percentage | # in Genome | *P* value |
|---|---|---|---|---|
| cell cycle | 47 | 0.032571033 | 395 | 2.60E-09 |
| microtubule-based process | 26 | 0.018018018 | 174 | 1.20E-07 |
| cell cycle process | 30 | 0.020790021 | 227 | 2.10E-07 |
| cell division | 24 | 0.016632017 | 165 | 6.00E-07 |
| mitotic cell cycle process | 20 | 0.013860014 | 122 | 7.60E-07 |
| microtubule-based movement | 17 | 0.011781012 | 93 | 1.10E-06 |
| DNA conformation change | 22 | 0.015246015 | 148 | 1.20E-06 |
| movement of cell or subcellular | 17 | 0.011781012 | 98 | 2.30E-06 |
| chromosome organization | 46 | 0.031878032 | 483 | 2.80E-06 |
| organelle organization | 90 | 0.062370062 | 1203 | 3.30E-06 |
| mitotic cell cycle | 23 | 0.015939016 | 173 | 5.00E-06 |
| cytokinesis | 10 | 0.006930007 | 45 | 3.10E-05 |
| cellular component organization | 130 | 0.09009009 | 2011 | 3.70E-05 |
| DNA replication initiation | 7 | 0.004851005 | 23 | 5.60E-05 |
| mitotic cytokinesis | 8 | 0.005544006 | 31 | 6.10E-05 |
| cytoskeleton-dependent cytokinesis | 8 | 0.005544006 | 32 | 7.90E-05 |
| positive regulation of histone methylation | 3 | 0.002079002 | 3 | 9.80E-05 |
| DNA geometric change | 12 | 0.008316008 | 77 | 0.0002 |
| DNA duplex unwinding | 12 | 0.008316008 | 77 | 0.0002 |
| response to water | 14 | 0.00970201 | 101 | 0.00022 |
| DNA replication | 16 | 0.011088011 | 127 | 0.00025 |
| DNA-dependent DNA replication | 12 | 0.008316008 | 79 | 0.00026 |
| positive regulation of chromosome | 4 | 0.002772003 | 8 | 0.00027 |
| cytokinetic process | 5 | 0.003465003 | 15 | 0.00042 |
| mitotic cytokinetic process | 5 | 0.003465003 | 15 | 0.00042 |
| cellular component organization or | 139 | 0.096327096 | 2309 | 0.00051 |
| response to water deprivation | 13 | 0.009009009 | 98 | 0.00057 |
| vesicle docking | 8 | 0.005544006 | 43 | 0.00069 |
| nuclear chromosome segregation | 11 | 0.007623008 | 76 | 0.00071 |

| | | | | |
|---|---|---|---|---|
| **sister chromatid segregation** | 9 | 0.006237006 | 54 | 0.00075 |
| **membrane docking** | 8 | 0.005544006 | 44 | 0.00081 |
| **organelle localization by membrane** | 8 | 0.005544006 | 44 | 0.00081 |
| **positive regulation of protein modification** | 11 | 0.007623008 | 79 | 0.00098 |
| **negative regulation of transcription, DNA-** | 10 | 0.006930007 | 68 | 0.00107 |
| **response to acid chemical** | 27 | 0.018711019 | 312 | 0.00135 |
| **chromosome segregation** | 11 | 0.007623008 | 83 | 0.00148 |
| **positive regulation of histone modification** | 3 | 0.002079002 | 6 | 0.00177 |
| **positive regulation of chromatin** | 3 | 0.002079002 | 6 | 0.00177 |
| **regulation of response to salt stress** | 5 | 0.003465003 | 20 | 0.00181 |
| **maintenance of chromatin silencing** | 2 | 0.001386001 | 2 | 0.00213 |
| **transposition, RNA-mediated** | 2 | 0.001386001 | 2 | 0.00213 |
| **maintenance of protein location in nucleus** | 2 | 0.001386001 | 2 | 0.00213 |
| **positive regulation of histone H3-K9** | 2 | 0.001386001 | 2 | 0.00213 |
| **regulation of jasmonic acid biosynthetic** | 2 | 0.001386001 | 2 | 0.00213 |
| **regulation of histone H4 acetylation** | 2 | 0.001386001 | 2 | 0.00213 |
| **negative regulation of histone H4** | 2 | 0.001386001 | 2 | 0.00213 |
| **DNA metabolic process** | 45 | 0.031185031 | 625 | 0.00218 |
| **regulation of response to osmotic stress** | 5 | 0.003465003 | 21 | 0.00228 |
| **cytokinesis by cell plate formation** | 4 | 0.002772003 | 13 | 0.00232 |
| **response to salt stress** | 14 | 0.00970201 | 129 | 0.00257 |
| **negative regulation of nucleobase-** | 12 | 0.008316008 | 102 | 0.0026 |
| **negative regulation of RNA biosynthetic** | 11 | 0.007623008 | 89 | 0.00262 |
| **negative regulation of nucleic acid-** | 11 | 0.007623008 | 89 | 0.00262 |
| **chaperone mediated protein folding** | 3 | 0.002079002 | 7 | 0.00299 |
| **negative regulation of RNA metabolic** | 11 | 0.007623008 | 92 | 0.0034 |
| **chromatin silencing** | 6 | 0.004158004 | 33 | 0.00364 |
| **cell wall biogenesis** | 13 | 0.009009009 | 121 | 0.00392 |
| **asymmetric cell division** | 3 | 0.002079002 | 8 | 0.00463 |
| **sodium ion import** | 3 | 0.002079002 | 8 | 0.00463 |
| **inorganic cation import across plasma** | 3 | 0.002079002 | 8 | 0.00463 |
| **sodium ion import across plasma** | 3 | 0.002079002 | 8 | 0.00463 |
| **inorganic ion import across plasma** | 3 | 0.002079002 | 8 | 0.00463 |
| **DNA packaging** | 8 | 0.005544006 | 59 | 0.00553 |
| **response to abiotic stimulus** | 42 | 0.029106029 | 604 | 0.00555 |
| **cell wall organization** | 29 | 0.02009702 | 380 | 0.00577 |
| **menaquinone metabolic process** | 2 | 0.001386001 | 3 | 0.00621 |
| **menaquinone biosynthetic process** | 2 | 0.001386001 | 3 | 0.00621 |
| **regulation of jasmonic acid metabolic** | 2 | 0.001386001 | 3 | 0.00621 |
| **bundle sheath cell fate specification** | 2 | 0.001386001 | 3 | 0.00621 |

| | | | | |
|---|---|---|---|---|
| **external encapsulating structure** | 30 | 0.020790021 | 399 | 0.00621 |
| **microsporogenesis** | 3 | 0.002079002 | 9 | 0.0067 |
| **positive regulation of protein** | 3 | 0.002079002 | 9 | 0.0067 |
| **'de novo' posttranslational protein folding** | 3 | 0.002079002 | 9 | 0.0067 |
| **positive regulation of ubiquitin-protein** | 3 | 0.002079002 | 9 | 0.0067 |
| **import across plasma membrane** | 3 | 0.002079002 | 9 | 0.0067 |
| **positive regulation of protein modification** | 3 | 0.002079002 | 9 | 0.0067 |
| **response to inorganic substance** | 20 | 0.013860014 | 236 | 0.00674 |
| **negative regulation of gene expression,** | 6 | 0.004158004 | 38 | 0.00745 |
| **response to osmotic stress** | 14 | 0.00970201 | 147 | 0.00822 |
| **positive regulation of response to salt** | 4 | 0.002772003 | 18 | 0.00825 |
| **regulation of mitotic cell cycle** | 10 | 0.006930007 | 90 | 0.00854 |
| **DNA repair** | 25 | 0.017325017 | 325 | 0.00901 |
| **maintenance of DNA methylation** | 3 | 0.002079002 | 10 | 0.00925 |
| **regulation of histone methylation** | 3 | 0.002079002 | 10 | 0.00925 |
| **organelle fusion** | 9 | 0.006237006 | 78 | 0.00969 |
| **positive regulation of protein metabolic** | 13 | 0.009009009 | 136 | 0.01033 |
| **mitotic sister chromatid segregation** | 6 | 0.004158004 | 41 | 0.0108 |
| **vesicle fusion** | 8 | 0.005544006 | 66 | 0.01081 |
| **response to oxygen-containing compound** | 31 | 0.021483021 | 435 | 0.01142 |
| **organelle membrane fusion** | 8 | 0.005544006 | 67 | 0.01179 |
| **allantoin catabolic process** | 2 | 0.001386001 | 4 | 0.01203 |
| **mitotic chromosome condensation** | 2 | 0.001386001 | 4 | 0.01203 |
| **chloroplast-nucleus signaling pathway** | 2 | 0.001386001 | 4 | 0.01203 |
| **cellular amide catabolic process** | 2 | 0.001386001 | 4 | 0.01203 |
| **histone H3-K9 methylation** | 2 | 0.001386001 | 4 | 0.01203 |
| **regulation of histone H3-K9 methylation** | 2 | 0.001386001 | 4 | 0.01203 |
| **histone H3-K9 modification** | 2 | 0.001386001 | 4 | 0.01203 |
| **response to chitin** | 3 | 0.002079002 | 11 | 0.01229 |
| **exocytosis** | 10 | 0.006930007 | 96 | 0.01318 |
| **DNA alkylation** | 4 | 0.002772003 | 21 | 0.01446 |
| **DNA methylation** | 4 | 0.002772003 | 21 | 0.01446 |
| **chromosome separation** | 5 | 0.003465003 | 32 | 0.01487 |
| **response to cold** | 10 | 0.006930007 | 98 | 0.01508 |
| **regulation of developmental growth** | 6 | 0.004158004 | 44 | 0.0151 |
| **cold acclimation** | 3 | 0.002079002 | 12 | 0.01583 |
| **jasmonic acid biosynthetic process** | 3 | 0.002079002 | 12 | 0.01583 |
| **cellulose microfibril organization** | 3 | 0.002079002 | 12 | 0.01583 |
| **sexual sporulation** | 3 | 0.002079002 | 12 | 0.01583 |
| **sporulation** | 3 | 0.002079002 | 12 | 0.01583 |

| | | | | |
|---|---|---|---|---|
| **plant-type sporogenesis** | 3 | 0.002079002 | 12 | 0.01583 |
| **regulation of ubiquitin-protein transferase** | 3 | 0.002079002 | 12 | 0.01583 |
| **cell wall assembly** | 3 | 0.002079002 | 12 | 0.01583 |
| **plant-type cell wall assembly** | 3 | 0.002079002 | 12 | 0.01583 |
| **meiotic cell cycle process** | 8 | 0.005544006 | 71 | 0.0164 |
| **meiotic cell cycle** | 9 | 0.006237006 | 85 | 0.01648 |
| **cellulose biosynthetic process** | 6 | 0.004158004 | 45 | 0.01676 |
| **regulation of cell cycle process** | 7 | 0.004851005 | 58 | 0.01701 |
| **cellular response to DNA damage stimulus** | 25 | 0.017325017 | 344 | 0.01736 |
| **organelle localization** | 10 | 0.006930007 | 101 | 0.01831 |
| **allantoin metabolic process** | 2 | 0.001386001 | 5 | 0.01944 |
| **methylation-dependent chromatin** | 2 | 0.001386001 | 5 | 0.01944 |
| **regulation of gene expression by genetic** | 2 | 0.001386001 | 5 | 0.01944 |
| **pyruvate transport** | 2 | 0.001386001 | 5 | 0.01944 |
| **mitochondrial pyruvate transmembrane** | 2 | 0.001386001 | 5 | 0.01944 |
| **cellularization** | 2 | 0.001386001 | 5 | 0.01944 |
| **radial pattern formation** | 2 | 0.001386001 | 5 | 0.01944 |
| **negative regulation of histone acetylation** | 2 | 0.001386001 | 5 | 0.01944 |
| **response to decreased oxygen levels** | 2 | 0.001386001 | 5 | 0.01944 |
| **regulation of fatty acid biosynthetic** | 2 | 0.001386001 | 5 | 0.01944 |
| **regulation of DNA methylation** | 2 | 0.001386001 | 5 | 0.01944 |
| **regulation of seed growth** | 2 | 0.001386001 | 5 | 0.01944 |
| **pyruvate transmembrane transport** | 2 | 0.001386001 | 5 | 0.01944 |
| **negative regulation of protein acetylation** | 2 | 0.001386001 | 5 | 0.01944 |
| **negative regulation of peptidyl-lysine** | 2 | 0.001386001 | 5 | 0.01944 |
| **branched-chain amino acid biosynthetic** | 4 | 0.002772003 | 23 | 0.0199 |
| **nucleosome assembly** | 6 | 0.004158004 | 47 | 0.02045 |
| **cellular amino acid catabolic process** | 6 | 0.004158004 | 47 | 0.02045 |
| **positive regulation of transferase activity** | 8 | 0.005544006 | 74 | 0.02061 |
| **cellular response to stress** | 39 | 0.027027027 | 602 | 0.02122 |
| **positive regulation of cellular protein** | 12 | 0.008316008 | 135 | 0.02259 |
| **positive regulation of organelle** | 4 | 0.002772003 | 24 | 0.02302 |
| **membrane fusion** | 8 | 0.005544006 | 76 | 0.02381 |
| **chromatin organization** | 21 | 0.014553015 | 285 | 0.02414 |
| **regulation of histone modification** | 3 | 0.002079002 | 14 | 0.02445 |
| **regulation of protein ubiquitination** | 3 | 0.002079002 | 14 | 0.02445 |
| **purine-containing compound catabolic** | 3 | 0.002079002 | 14 | 0.02445 |
| **regulation of protein modification by** | 3 | 0.002079002 | 14 | 0.02445 |
| **mitochondrial transmembrane transport** | 3 | 0.002079002 | 14 | 0.02445 |
| **secretion by cell** | 10 | 0.006930007 | 106 | 0.02477 |

| | | | | |
|---|---|---|---|---|
| **response to chemical** | 68 | 0.047124047 | 1161 | 0.02535 |
| **response to organic substance** | 41 | 0.028413028 | 647 | 0.02543 |
| **beta-glucan biosynthetic process** | 7 | 0.004851005 | 63 | 0.02572 |
| **nuclear division** | 10 | 0.006930007 | 107 | 0.02624 |
| **response to salicylic acid** | 4 | 0.002772003 | 25 | 0.02643 |
| **xyloglucan metabolic process** | 6 | 0.004158004 | 50 | 0.02697 |
| **purine nucleobase catabolic process** | 2 | 0.001386001 | 6 | 0.02828 |
| **nucleoside triphosphate catabolic process** | 2 | 0.001386001 | 6 | 0.02828 |
| **adaxial/abaxial pattern specification** | 2 | 0.001386001 | 6 | 0.02828 |
| **root meristem growth** | 2 | 0.001386001 | 6 | 0.02828 |
| **gas transport** | 2 | 0.001386001 | 6 | 0.02828 |
| **oxygen transport** | 2 | 0.001386001 | 6 | 0.02828 |
| **regulation of exocytosis** | 2 | 0.001386001 | 6 | 0.02828 |
| **regulation of fatty acid metabolic process** | 2 | 0.001386001 | 6 | 0.02828 |
| **chromosome condensation** | 2 | 0.001386001 | 6 | 0.02828 |
| **mitochondrial respiratory chain complex** | 2 | 0.001386001 | 6 | 0.02828 |
| **regulation of histone acetylation** | 2 | 0.001386001 | 6 | 0.02828 |
| **histone H4 acetylation** | 2 | 0.001386001 | 6 | 0.02828 |
| **response to freezing** | 2 | 0.001386001 | 6 | 0.02828 |
| **regulation of secretion** | 2 | 0.001386001 | 6 | 0.02828 |
| **regulation of mitotic spindle organization** | 2 | 0.001386001 | 6 | 0.02828 |
| **response to oxygen levels** | 2 | 0.001386001 | 6 | 0.02828 |
| **genetic imprinting** | 2 | 0.001386001 | 6 | 0.02828 |
| **seed growth** | 2 | 0.001386001 | 6 | 0.02828 |
| **regulation of chlorophyll metabolic** | 2 | 0.001386001 | 6 | 0.02828 |
| **regulation of spindle organization** | 2 | 0.001386001 | 6 | 0.02828 |
| **regulation of protein acetylation** | 2 | 0.001386001 | 6 | 0.02828 |
| **regulation of secretion by cell** | 2 | 0.001386001 | 6 | 0.02828 |
| **positive regulation of ubiquitin protein** | 2 | 0.001386001 | 6 | 0.02828 |
| **regulation of peptidyl-lysine acetylation** | 2 | 0.001386001 | 6 | 0.02828 |
| **organelle fission** | 12 | 0.008316008 | 140 | 0.02899 |
| **postreplication repair** | 3 | 0.002079002 | 15 | 0.02954 |
| **regulation of lipid biosynthetic process** | 3 | 0.002079002 | 15 | 0.02954 |
| **mitotic cell cycle phase transition** | 5 | 0.003465003 | 38 | 0.02959 |
| **response to temperature stimulus** | 14 | 0.00970201 | 173 | 0.02995 |
| **megagametogenesis** | 4 | 0.002772003 | 26 | 0.03013 |
| **cell cycle phase transition** | 5 | 0.003465003 | 39 | 0.03269 |
| **cytoskeleton organization** | 13 | 0.009009009 | 159 | 0.03312 |
| **cell wall organization or biogenesis** | 34 | 0.023562024 | 530 | 0.03381 |
| **DNA methylation or demethylation** | 4 | 0.002772003 | 27 | 0.03412 |

| | | | | |
|---|---|---|---|---|
| **secretion** | 10 | 0.006930007 | 112 | 0.03452 |
| **microtubule cytoskeleton organization** | 7 | 0.004851005 | 67 | 0.03458 |
| **regulation of protein modification process** | 13 | 0.009009009 | 160 | 0.03459 |
| **vacuolar transport** | 6 | 0.004158004 | 53 | 0.03473 |
| **chromatin assembly** | 6 | 0.004158004 | 53 | 0.03473 |
| **nucleosome organization** | 6 | 0.004158004 | 53 | 0.03473 |
| **mitotic nuclear division** | 6 | 0.004158004 | 53 | 0.03473 |
| **response to lipid** | 17 | 0.011781012 | 227 | 0.03473 |
| **plant-type primary cell wall biogenesis** | 3 | 0.002079002 | 16 | 0.03513 |
| **negative regulation of nitrogen compound** | 20 | 0.013860014 | 280 | 0.03614 |
| **G1/S transition of mitotic cell cycle** | 2 | 0.001386001 | 7 | 0.03839 |
| **glycine catabolic process** | 2 | 0.001386001 | 7 | 0.03839 |
| **deoxyribonucleotide biosynthetic process** | 2 | 0.001386001 | 7 | 0.03839 |
| **genetic transfer** | 2 | 0.001386001 | 7 | 0.03839 |
| **DNA mediated transformation** | 2 | 0.001386001 | 7 | 0.03839 |
| **poly(A)+ mRNA export from nucleus** | 2 | 0.001386001 | 7 | 0.03839 |
| **negative regulation of histone** | 2 | 0.001386001 | 7 | 0.03839 |
| **UDP-L-arabinose metabolic process** | 2 | 0.001386001 | 7 | 0.03839 |
| **multi-organism cellular process** | 2 | 0.001386001 | 7 | 0.03839 |
| **cell cycle G1/S phase transition** | 2 | 0.001386001 | 7 | 0.03839 |
| **regulation of microtubule cytoskeleton** | 2 | 0.001386001 | 7 | 0.03839 |
| **mitochondrial respiratory chain complex** | 2 | 0.001386001 | 7 | 0.03839 |
| **nucleoside phosphate catabolic process** | 2 | 0.001386001 | 7 | 0.03839 |
| **branched-chain amino acid metabolic** | 4 | 0.002772003 | 28 | 0.0384 |
| **response to topologically incorrect protein** | 4 | 0.002772003 | 28 | 0.0384 |
| **response to stimulus** | 175 | 0.121275121 | 3356 | 0.03876 |
| **vesicle organization** | 8 | 0.005544006 | 84 | 0.0401 |
| **jasmonic acid metabolic process** | 3 | 0.002079002 | 17 | 0.04123 |
| **cellular glucan metabolic process** | 15 | 0.01039501 | 198 | 0.04168 |
| **DNA modification** | 4 | 0.002772003 | 29 | 0.04297 |
| **cellular polysaccharide metabolic process** | 17 | 0.011781012 | 235 | 0.04573 |
| **activation of MAPK activity involved in** | 1 | 0.000693001 | 1 | 0.04623 |
| **photoreactive repair** | 1 | 0.000693001 | 1 | 0.04623 |
| **assembly of actomyosin apparatus** | 1 | 0.000693001 | 1 | 0.04623 |
| **phragmoplast assembly** | 1 | 0.000693001 | 1 | 0.04623 |
| **action potential** | 1 | 0.000693001 | 1 | 0.04623 |
| **dUMP biosynthetic process** | 1 | 0.000693001 | 1 | 0.04623 |
| **pyrimidine nucleotide catabolic process** | 1 | 0.000693001 | 1 | 0.04623 |
| **DNA unwinding involved in DNA** | 1 | 0.000693001 | 1 | 0.04623 |
| **osmosensory signaling pathway** | 1 | 0.000693001 | 1 | 0.04623 |

| | | | | |
|---|---|---|---|---|
| **pyrimidine nucleoside triphosphate** | 1 | 0.000693001 | 1 | 0.04623 |
| **pyrimidine deoxyribonucleoside** | 1 | 0.000693001 | 1 | 0.04623 |
| **pyrimidine deoxyribonucleotide catabolic** | 1 | 0.000693001 | 1 | 0.04623 |
| **radial microtubular system formation** | 1 | 0.000693001 | 1 | 0.04623 |
| **endosperm cellularization** | 1 | 0.000693001 | 1 | 0.04623 |
| **vegetative meristem growth** | 1 | 0.000693001 | 1 | 0.04623 |
| **magnesium ion homeostasis** | 1 | 0.000693001 | 1 | 0.04623 |
| **mitochondrial DNA metabolic process** | 1 | 0.000693001 | 1 | 0.04623 |
| **chlorophyll cycle** | 1 | 0.000693001 | 1 | 0.04623 |
| **DNA rewinding** | 1 | 0.000693001 | 1 | 0.04623 |
| **dUMP metabolic process** | 1 | 0.000693001 | 1 | 0.04623 |
| **dUTP metabolic process** | 1 | 0.000693001 | 1 | 0.04623 |
| **dUTP catabolic process** | 1 | 0.000693001 | 1 | 0.04623 |
| **membrane depolarization** | 1 | 0.000693001 | 1 | 0.04623 |
| **positive regulation of histone H3-K27** | 1 | 0.000693001 | 1 | 0.04623 |
| **pri-miRNA transcription from RNA** | 1 | 0.000693001 | 1 | 0.04623 |
| **membrane depolarization during action** | 1 | 0.000693001 | 1 | 0.04623 |
| **tubulin deacetylation** | 1 | 0.000693001 | 1 | 0.04623 |
| **nicotinate metabolic process** | 1 | 0.000693001 | 1 | 0.04623 |
| **assembly of actomyosin apparatus** | 1 | 0.000693001 | 1 | 0.04623 |
| **positive regulation of defense response to** | 1 | 0.000693001 | 1 | 0.04623 |
| **polyuridylation-dependent mRNA** | 1 | 0.000693001 | 1 | 0.04623 |
| **malonyl-CoA biosynthetic process** | 1 | 0.000693001 | 1 | 0.04623 |
| **galactose metabolic process** | 3 | 0.002079002 | 18 | 0.04783 |
| **purine nucleobase metabolic process** | 3 | 0.002079002 | 18 | 0.04783 |
| **maintenance of location in cell** | 3 | 0.002079002 | 18 | 0.04783 |
| **G-protein coupled receptor signaling** | 4 | 0.002772003 | 30 | 0.04783 |
| **cellulose metabolic process** | 7 | 0.004851005 | 72 | 0.04823 |
| **gametophyte development** | 10 | 0.006930007 | 119 | 0.04894 |
| **membrane organization** | 12 | 0.008316008 | 152 | 0.04948 |
| **respiratory chain complex IV assembly** | 2 | 0.001386001 | 8 | 0.04964 |
| **serine family amino acid catabolic process** | 2 | 0.001386001 | 8 | 0.04964 |
| **nucleoside catabolic process** | 2 | 0.001386001 | 8 | 0.04964 |
| **DNA methylation on cytosine** | 2 | 0.001386001 | 8 | 0.04964 |
| **pyrimidine-containing compound** | 2 | 0.001386001 | 8 | 0.04964 |
| **maintenance of protein localization in** | 2 | 0.001386001 | 8 | 0.04964 |
| **C-5 methylation of cytosine** | 2 | 0.001386001 | 8 | 0.04964 |
| **regulation of tetrapyrrole metabolic** | 2 | 0.001386001 | 8 | 0.04964 |
| **glycosyl compound catabolic process** | 2 | 0.001386001 | 8 | 0.04964 |

**Supplementary Table 9. Gene ontology enrichment of the 268 genes differentially expressed between the *Pi-ta*$^{-}$ (X2) and *Pi-ta*$^{0}$ (X0) lines.** *Pi-ta*$^{-}$ and *Pi-ta*$^{0}$ lines are the *Pi-ta* knockout lines and their sibling line without mutations in *Pi-ta* respectively. GO annotations are from experimentally verified datasets and differentially expressed genes were compared to the entire annotated *Oryza sativa* genome. Percentage is the number of genes enriched in one GO pathway divided the total number of genes differentially expressed between the *Pi-ta*$^{-}$ and *Pi-ta*$^{WT}$ lines.

| GO annotation | # DE in *Pi-ta*$^{-}$ vs *Pi-ta*$^{0}$ | Percentage | # in Genome | *P* value |
|---|---|---|---|---|
| **oxidation-reduction process** | 12 | 0.12 | 1861 | 0.0278 |
| **carbohydrate metabolic process** | 8 | 0.08 | 949 | 0.0181 |
| **integrin-mediated signaling pathway** | 1 | 0.01 | 1 | 0.0036 |
| **mature ribosome assembly** | 1 | 0.01 | 5 | 0.0176 |
| **assembly of large subunit precursor of preribosome** | 1 | 0.01 | 7 | 0.0246 |
| **mRNA cleavage involved in mRNA processing** | 1 | 0.01 | 5 | 0.0176 |
| **pre-mRNA cleavage required for polyadenylation** | 1 | 0.01 | 5 | 0.0176 |
| **mRNA cleavage** | 1 | 0.01 | 10 | 0.035 |
| **establishment of sister chromatid cohesion** | 1 | 0.01 | 3 | 0.0106 |
| **viral process** | 1 | 0.01 | 13 | 0.0452 |
| **modification of morphology or physiology of other organism involved in symbiotic interaction** | 1 | 0.01 | 8 | 0.0281 |
| **modification of morphology or physiology of other organism** | 1 | 0.01 | 8 | 0.0281 |
| **interaction with host** | 1 | 0.01 | 12 | 0.0418 |
| **modulation by virus of host morphology or physiology** | 1 | 0.01 | 4 | 0.0141 |
| **modification by symbiont of host morphology or physiology** | 1 | 0.01 | 8 | 0.0281 |
| **oxidative photosynthetic carbon pathway** | 1 | 0.01 | 5 | 0.0176 |
| **polyphosphate metabolic process** | 1 | 0.01 | 2 | 0.0071 |
| **polyphosphate biosynthetic process** | 1 | 0.01 | 1 | 0.0036 |
| **phosphate ion transmembrane transport** | 1 | 0.01 | 13 | 0.0452 |
| **inositol phosphate-mediated signaling** | 1 | 0.01 | 1 | 0.0036 |
| **UTP metabolic process** | 1 | 0.01 | 5 | 0.0176 |
| **UTP biosynthetic process** | 1 | 0.01 | 5 | 0.0176 |
| **guanosine-containing compound biosynthetic** | 1 | 0.01 | 9 | 0.0315 |

| | | | | |
|---|---|---|---|---|
| **process** | | | | |
| **GTP metabolic process** | 1 | 0.01 | 9 | 0.0315 |
| **GTP biosynthetic process** | 1 | 0.01 | 6 | 0.0211 |
| **hormone metabolic process** | 2 | 0.02 | 86 | 0.0374 |
| **hormone biosynthetic process** | 2 | 0.02 | 66 | 0.023 |
| **auxin metabolic process** | 1 | 0.01 | 13 | 0.0452 |
| **auxin biosynthetic process** | 1 | 0.01 | 9 | 0.0315 |
| **leaf development** | 2 | 0.02 | 78 | 0.0313 |
| **negative gravitropism** | 1 | 0.01 | 11 | 0.0384 |
